# Attention selectively reshapes the geometry of distributed semantic representation

**DOI:** 10.1101/045252

**Authors:** Samuel A. Nastase, Andrew C. Connolly, Nikolaas N. Oosterhof, Yaroslav O. Halchenko, J. Swaroop Guntupalli, Matteo Visconti di Oleggio Castello, Jason Gors, M. Ida Gobbini, James V. Haxby

## Abstract

Humans prioritize different semantic qualities of a complex stimulus depending on their behavioral goals. These semantic features are encoded in distributed neural populations, yet it is unclear how attention might operate across these distributed representations. To address this, we presented participants with naturalistic video clips of animals behaving in their natural environments while the participants attended to either behavior or taxonomy. We used models of representational geometry to investigate how attentional allocation affects the distributed neural representation of animal behavior and taxonomy. Attending to animal behavior transiently increased the discriminability of distributed population codes for observed actions in anterior intraparietal, pericentral, and ventral temporal cortices. Attending to animal taxonomy while viewing the same stimuli increased the discriminability of distributed animal category representations in ventral temporal cortex. For both tasks, attention selectively enhanced the discriminability of response patterns along behaviorally relevant dimensions. These findings suggest that behavioral goals alter how the brain extracts semantic features from the visual world. Attention effectively disentangles population responses for downstream read-out by sculpting representational geometry in late-stage perceptual areas.

## Introduction

The brain’s information processing machinery operates dynamically to accommodate diverse behavioral goals. Selective attention reduces the complexity of information processing by prioritizing representational content relevant to the task at hand (Tsotsos 2011). The attention literature has focused mostly on early vision, employing rudimentary visual stimuli and simple tasks to probe task-related changes in the representation of low-level visual information, such as orientation and motion direction (Carrasco 2011). Humans, however, perceive and act on the world in terms of both semantically-rich representations and complex behavioral goals. Naturalistic stimuli, although less controlled, serve to convey richer perceptual and semantic information, and have been shown to reliably drive neural responses (Hasson et al. 2004; Haxby et al. 2011; Huth et al. 2012, 2016; Guntupalli et al. 2016).

The brain encodes this sort of complex information in high-dimensional representational spaces grounded in the concerted activity of distributed populations of neurons (Averbeck et al. 2006; Kriegeskorte et al. 2008a; Haxby et al. 2014). Population coding is an important motif in neural information processing across species (Dayan and Abbott 2001), and has been well-characterized in early vision (Chen et al. 2006; Miyawaki et al. 2008; Graf et al. 2011), face and object recognition (Rolls and Tovee 1995; Hung et al. 2005; Kiani et al. 2007; Freiwald and Tsao 2010), and other sensorimotor and cognitive domains (Georgopoulos et al. 1986; Lewis and Kristan 1998; Uchida et al. 2000; Rigotti et al. 2013). Multivariate decoding analyses of human neuroimaging data have allowed us to leverage distributed patterns of cortical activation to provide a window into the representation of high-level semantic information (Haxby et al. 2001, 2014; Kriegeskorte et al. 2008a; Mitchell et al. 2008; Oosterhof et al. 2010, 2012; Connolly et al. 2012, 2016; Huth et al. 2012; Sha et al. 2015), but these studies generally assume that neural representations are relatively stable, rather than dynamic or context-dependent.

Electrophysiological work on attentional modulation has typically been constrained to single neurons (Treue and Martínez Trujillo 1999; Reynolds et al. 2000; Reynolds and Heeger 2009). For example, attention shifts the balance between excitatory and suppressive neural activity to accentuate the responses of neurons tuned to task-relevant features (Reynolds and Heeger 2009), and object categorization training increases selectivity for task-relevant stimulus features in in macaque temporal cortex neurons (Sigala and Logothetis 2002). More recent work has suggested that task demands may alter population encoding to sharpen attended representations (Cohen and Maunsell 2009; Ruff and Cohen 2014; Downer et al. 2015). In line with this, a handful of recent neuroimaging studies have examined how task demands affect multivariate pattern classification (Serences and Boynton 2007; Peelen et al. 2009; Jehee et al. 2011; Brouwer and Heeger 2013; Sprague and Serences 2013; Harel et al. 2014; Erez and Duncan 2015). In particular, Brouwer and Heeger (2013) demonstrated that when participants perform a color naming task, distributed neural representations of color in two early visual areas become more categorical—that is, the neural color space is altered such that within-category distances decrease while between-category colors increase. In a related approach, Çukur and colleagues (2013) used a natural vision paradigm to demonstrate that performing a covert visual search task for either humans or vehicles in natural scenes drives widespread shifts in voxelwise semantic tuning, even when these target objects are not present in the stimulus. With the exception of this study, most prior work has investigated only simple visual stimuli such as oriented gratings, moving dots, colors, and static object images. The current study aims to directly investigate task-related changes in the geometry of distributed neural representation of high-level visual and semantic information about animal taxonomy and behavior conveyed by dynamic, naturalistic stimuli.

We hypothesized that, in order to interface with distributed neural representations, attention may operate in a distributed fashion as well—that is, by selectively reshaping representational geometry (Edelman 1998; Kriegeskorte and Kievit 2013). This hypothesis was motivated by behavioral and theoretical work suggesting that attention may facilitate categorization by expanding psychological distances along task-relevant stimulus dimensions and collapsing task-irrelevant distinctions (Nosofsky 1986; Kruschke 1992). Here we aimed to provide neural evidence for this phenomenon by examining how task demands affect the distributed neural representation of two types of semantic information thought to rely on distributed population codes: animal taxonomy (Connolly et al. 2012, 2016; Sha et al. 2015) and behavior (Oosterhof et al. 2010, 2012, 2013). We operationalize attention broadly in this context as the modulatory effect of top-down task demands on stimulus-evoked neural representation; at minimum, the 1-back task requires participants to categorize stimuli, maintain the previously observed category in working memory, and compare the currently observed category with the prior category, and execute (or withhold) a motor response. To expand on previous work, we used dynamic, naturalistic video clips of animals behaving in their natural environments. These stimuli not only convey information about animal form or category, but also behavior, allowing us to examine how attention affects the neural representation of observed actions (Oosterhof et al. 2013), which has not previously been studied. Categorical models of representational geometry were employed to demonstrate that attention selectively alters distances between neural representations of both animal taxonomy and behavior along task-relevant dimensions.

## Materials and Methods

### Participants

Twelve right-handed adults (seven female; mean age = 25.4 ± 2.6 SD years) with normal or corrected-to-normal vision participated in the attention experiment. Participants reported no neurological conditions. Additionally, 19 adults, including the 12 from the attention experiment, participated in a separate scanning session for the purposes of hyperalignment. All participants gave written, informed consent prior to participating in the study, and the study was approved by the Institutional Review Board of Dartmouth College.

### Stimuli and design

Each of the 20 conditions in the fully-crossed design comprised two unique exemplar clips of animals from five taxonomic categories (primates, ungulates, birds, reptiles, insects) performing actions from four behavioral categories (eating, fighting, running, swimming) as well as their horizontally-flipped counterparts, for a total of 40 clips and 80 total exemplars (Supplementary Table 1, Supplementary Video 1). The four behavioral categories and five taxonomic categories roughly correspond to intermediate levels of noun and verb category hierarchies (Fellbaum 1990; Rosch 1978). Note that although the taxonomy and behavior factors are orthogonalized at the level of categorization relevant for the task, orthogonalizing lower-level variables (e.g., the specific animal performing each behavior) is not feasible in naturalistic contexts. Each trial consisted of a 2 s video clip presented without sound followed by a 2 s fixation period in a rapid event-related design. Clips for the attention experiment were extracted from nature documentaries (*Life*, *Life of Mammals*, *Microcosmos*, *Planet Earth*) and YouTube videos matched for resolution. The clips used in the attention experiment were not included in the segment of the documentary presented for the purpose of hyperalignment. All 80 stimuli, as well as four behavior repetition events, four taxon repetition events, and four null events were presented in pseudorandom order in each of 10 runs, resulting in 92 events per run, plus 12 s fixation before and after the events of interest, for a total run length of 392 s (∼6.5 min). Ten unique runs were constructed for a total scan time of approximately 65 min, and run order was counterbalanced across participants. At the beginning of each run, participants were instructed to pay attention to either taxonomy or behavior types and press the button only when they observed a category repetition of that type. Prior to scanning, participants were verbally familiarized with the categories and their constituents (e.g., the “ungulates” category includes quadrupedal, hoofed, herbivorous mammals such as horses). There were five behavior attention runs and five taxonomy attention runs presented in counterbalanced order across participants.

For each run, a pseudorandom trial order was first constructed such that no taxonomic or behavioral categories were repeated (adjacent in the trial order). Next, four taxonomic category repetition events and four behavioral category repetition events were inserted as sparse catch trials such that a repetition event of each type fell on a random trial within each quarter of the run (without inducing unplanned repetitions). Each repetition event repeated either the taxonomic or behavioral category of the preceding stimulus and varied on the other dimension. There were no repetitions of the same clip exemplar (or its horizontal mirror image). Four additional 2 s null events consisting of only a fixation cross were inserted into each run to effect temporal jittering.

The same button was pressed for repetitions of both types. Button presses were only elicited by repetition events and were therefore sparse. Participants were informed that repetition events would be sparse and that they should not pay attention to or press the button if they noticed repetitions of the unattended type. Participants were only instructed to maintain fixation when the fixation cross was present, not during the presentation of the clips.

In an independent scanning session, participants were presented with approximately 63 min of the *Life* nature documentary narrated by David Attenborough for the purpose of hyperalignment. The documentary was presented in four runs of similar duration, and included both the visual and auditory tracks. In the movie session, participants were instructed to remain still and watch the documentary as though they were watching a movie at home. All stimuli were presented using PsychoPy (Peirce 2007).

### Image acquisition

All functional and structural images were acquired using a 3 T Philips Intera Achieva MRI scanner (Philips Medical Systems, Bothell, WA) with a 32-channel phased-array SENSE (SENSitivity Encoding) head coil. For the attention experiment, functional images were acquired in an interleaved fashion using single-shot gradient-echo echo-planar imaging with a SENSE reduction factor of 2 (TR/TE = 2000/35 ms, flip angle = 90°, resolution = 3 mm isotropic, matrix size = 80 × 80, FOV = 240 × 240 mm, 42 transverse slices with full brain coverage and no gap). Each run began with two dummy scans to allow for signal stabilization. For each participant 10 runs were collected, each consisting of 196 dynamic scans totaling 392 s (∼6.5 min). At the end of each scanning session, a structural scan was obtained using a high-resolution T1-weighted 3D turbo field echo sequence (TR/TE = 8.2/3.7 ms, flip angle = 8°, resolution = 1 mm isotropic, matrix size = 256 × 256 × 220, FOV = 240 × 188 × 220 mm).

For the movie session, functional images also were acquired in an interleaved order using single-shot gradient-echo echo-planar imaging (TR/TE = 2500/35 ms, flip angle = 90°, resolution = 3 mm isotropic, matrix size = 80 × 80, and FOV = 240 × 240 mm; 42 transverse slices with full brain coverage and no gap). Four runs were collected for each participant, consisting of 374, 346, 377, and 412 dynamic scans, totaling 935 s (∼15.6 min), 865 s (∼14.4 min), 942.5 s (∼15.7 min), and 1030 s (∼17.2 min), respectively. At the end of this session, a structural scan was obtained using a high-resolution T1-weighted 3D turbo field echo sequence (TR/TE = 8.2/3.7 ms, flip angle = 8°, resolution = 1 mm isotropic, matrix size = 256 × 256 × 220, and FOV = 240 × 188 × 220). For participants included in both the attention experiment and the movie session, structural images were registered and averaged to increase signal-to-noise ratio.

### Preprocessing

For each participant, functional time series data were de-spiked, corrected for slice timing and head motion, normalized to the ICBM 452 template in MNI space, and spatially smoothed with a 4 mm FWHM Gaussian kernel using AFNI (Cox 1996). Functional images were then motion-corrected in two passes: first, functional images were initially motion corrected, then averaged across time to create a high-contrast reference volume; motion correction parameters were then re-estimated in a second pass using the reference volume as the target. Affine transformation parameters were then estimated to coregister the reference volume and the participant’s averaged structural scans. Each participant’s averaged structural scan was then normalized to the ICBM 452 template in MNI space. These transformation matrices were concatenated and each participant’s data were motion-corrected and normalized to the template via the participant’s anatomical scan in a single interpolation step. All subsequent analyses were performed in MNI space. Signal intensities were normalized to percent signal change prior to applying the general linear model.

Functional time series from the *Life* movie session were analyzed using the same preprocessing pipeline. Prior to hyperalignment, time series data were bandpass filtered to remove frequencies higher than 0.1 Hz and lower than 0.0067 Hz. Head motion parameters and the mean time series derived from the FreeSurfer segmentation of the ventricles were regressed out of the signal.

Cortical surfaces were reconstructed from structural scans using FreeSurfer, aligned according to curvature patterns on the spherical surface projection (Fischl et al. 1999), and visualized using SUMA (Saad et al. 2004).

### General linear model

A general linear model (GLM) was used to estimate BOLD responses for each of the 20 conditions for each task using AFNI’s 3dREMLfit. Stimulus-evoked BOLD responses to each event were modeled using a simple hemodynamic response function (AFNI’s GAM response model) adjusted for a 2 s stimulus duration. Nuisance regressors accounting for repetition events, button presses, and head motion were included in the model. For representational similarity analyses, beta parameters were estimated over the five taxonomy attention runs, then separately over the five behavior attention runs. Time points subtending abrupt head movements greater than roughly 1 mm of displacement or 1 degree of rotation were censored when fitting the general linear model. Response patterns were estimated from the 80 trials in each run, excluding repetition trials. For each of the two attention conditions, four trials per run from each of five runs contributed to the stimulus-evoked response pattern for each taxonomic–behavioral category condition, meaning that each pattern was estimated from 20 trials presented in pseudorandom order over the course of five separate runs (interspersed with runs from the other attention condition). Therefore we expect these response patterns (and the subsequent neural representational geometries) to be relatively robust to instrumental noise, temporal autocorrelation and intrinsic physiological correlations in the preprocessed time series data (Henriksson et al. 2015). Betas for each voxel were z-scored across the 20 conditions per feature before and after hyperalignment, and prior to any multivariate analysis. Note that constructing neural representational dissimilarity matrices (RDMs) by computing the correlation distance between response pattern vectors (rather than, e.g., Euclidean distance) entails that the subsequent multivariate analyses are invariant to differences in regional-average activity levels within a searchlight or ROI (Kriegeskorte et al. 2008b). For searchlight classification analyses (Supplementary Fig. 2), beta parameters were estimated separately for each run.

### Whole-brain hyperalignment

Surface-based searchlight whole-brain hyperalignment (Haxby et al. 2011; Guntupalli et al. 2016) was performed based on data collected while participants viewed the *Life* nature documentary. Each surface-based searchlight referenced the 200 nearest voxels from the associated volume, selected based on their geodesic proximity to the searchlight center. The time series of response patterns elicited by the movie stimulus was rotated via the Procrustes transformation in order to achieve optimal functional alignment across participants and the estimated transformation matrices for each searchlight were aggregated (Supplementary Fig. 1*A*). Hyperalignment transformation parameters estimated from the movie data were then applied to the independent attention experiment data. Subsequent analyses were applied to the hyperaligned data. All multivariate pattern analyses were performed using the PyMVPA package (www.pymvpa.org; Hanke et al. 2009).

### Searchlight representational similarity regression

Representational similarity analysis (Kriegeskorte et al. 2008b) was applied using 100-voxel surface-based searchlights (Kriegeskorte et al. 2006; Oosterhof et al. 2011). Each surface-based searchlight referenced the 100 nearest voxels to the searchlight center based on geodesic distance on the cortical surface. Pairwise correlation distances between stimulus-evoked response patterns for the 20 conditions were computed separately for each task. These pairwise distances were collated into a representational dissimilarity matrix (RDM) describing the representational geometry for a patch of cortex (Kriegeskorte and Kievit 2013).

Two categorical target RDMs were constructed based on the experimental design: one of these RDMs discriminated the animal taxa invariant to behavior, the other discriminated the behaviors invariant to taxonomy. Least squares multiple regression was then used to model the observed neural RDM as a weighted sum of the two categorical target RDMs. For each searchlight, both the observed neural RDM and the target RDMs were ranked and standardized prior to regression (see Saltelli et al. 2004). Since we suspect the neural representational space does not respect the magnitude of dissimilarity specified by our models, we relax the linear constraint and ensure only monotonicity (analogous to Spearman correlation, keeping with Kriegeskorte et al. 2008b). Although applying the rank transform prior to least squares linear regression is relatively common practice, this approach may emphasize main effects at the expense of interaction effects; however, in the current experiment, we have no a priori hypotheses corresponding to interaction terms. Intercept terms in the estimated models were negligible across all searchlights, task conditions, and participants. The searchlight analysis was performed in the hyperaligned space, then the results were projected onto the cortical surface reconstruction for the participant serving as the reference participant in the hyperalignment algorithm.

### Statistical assessment of searchlight analysis

To assess the statistical significance of searchlight maps across participants, all maps were corrected for multiple comparisons without choosing an arbitrary uncorrected threshold using threshold-free cluster enhancement (TFCE) with the recommended values (Smith and Nichols 2009). A Monte Carlo simulation permuting condition labels was used to estimate a null TFCE distribution (Oosterhof et al. 2012, 2016). To test the null hypothesis that response patterns contain no information about the 20 taxonomy–behavior conditions separately for each task condition, we permuted the 20 condition labels. To test the null hypothesis that task does not affect representational geometry, we permuted the sign of the difference between regression coefficients (following representational similarity regression) for the two tasks. First, 100 null searchlight maps were generated for each participant by randomly permuting the 20 condition labels within each observed searchlight RDM, then computing the regression described above. Next, we randomly sampled (with replacement) from these null searchlight maps, computed the mean searchlight regression coefficient across participants for each random sample of 12 data sets (one from each participant), then computed TFCE. This resulted in a null TFCE map. We then repeated this resampling procedure 10,000 times to construct a null distribution of TFCE maps (Stelzer et al., 2013). The resulting group searchlight maps are thresholded at cluster-level *p* = .05 corrected for familywise error using TFCE, and the average regression coefficient across participants is plotted for surviving searchlights.

In the case of searchlight classification (Supplementary Fig. 2), labels were shuffled within each run and each category of the crossed factor (e.g., the four behavior labels were permuted within each of the five taxa), then the full cross-validation scheme was applied (Nastase et al. 2016). The resulting maps are similarly thresholded, with the average classification accuracy across participants plotted for surviving searchlights. For difference maps (Supplementary Fig. 3), clusters surviving correction for multiple comparisons are indicated by white contours and subthreshold searchlights are displayed transparently. This method for multiple comparisons correction was implemented using the CoSMoMVPA software package (cosmomvpa.org; Oosterhof et al. 2016).

To assess more global effects, task-related differences in regression coefficients across searchlights were computed separately for each categorical target RDM. We assessed whether the attentional task altered mean regression coefficients for both categorical target RDMs within searchlights containing information about behavioral and taxonomic categories. For the behavioral category target RDM, the mean regression coefficients were computed across all searchlight regression coefficients surviving statistical thresholding using TFCE (cluster-level *p* < .05) in either attention condition. A nonparametric randomization test was used to evaluate the significance of a task difference in the regression coefficient across participants. The sign of the attentional difference in the mean regression coefficient across searchlights was permuted across participants. This tests the null hypothesis that there is no systematic attentional difference in searchlight regression coefficients across participants. This procedure was repeated for the taxonomic category target RDM considering all searchlight regression coefficients that survived TFCE in both tasks.

### Identifying regions of interest

Cluster analysis was used to identify regions of the cortical surface characterized by shared representational geometry in an unsupervised manner (Connolly et al. 2012). Prior to cluster analysis, the observed neural RDMs for each surface-based searchlight were converted from correlation distances to Fisher transformed correlation values and averaged across participants. Gaussian mixture models were used to cluster searchlights according to their representational geometry at varying values of *k* components (clusters). Gaussian mixture modeling is a probabilistic generalization of the *k*-means algorithm, and models the 20,484 searchlights as a mixture of *k* overlapping Gaussian distributions in a 190-dimensional feature space defined by the upper triangular of the 20 × 20 observed neural RDM. The clustering algorithm was implemented using the scikit-learn machine learning library for Python (Pedregosa et al. 2011).

We evaluated the reproducibility of parcellations across participants at values of *k* from 2 to 30 using a split--half resampling approach (100 iterations per *k*) that has previously been applied to functional parcellations based on resting-state functional connectivity (Yeo et al. 2011). For each of 100 resampling iterations, half of the participants were randomly assigned to a training set, while the other half were assigned to a test set (Lange et al. 2004). Surface-based searchlight RDMs for each participant were then averaged across participants within the separate training and test sets. Gaussian mixture models were estimated on the training set for each of *k* components ranging from 2 to 30. Test data were then assigned to the nearest cluster mean of the model estimated from the training data. A separate mixture model was then estimated for the test data, and the predicted cluster labels (based on the training data) were compared to the actual cluster labels using adjusted mutual information (AMI; Thirion et al. 2014). AMI compares cluster solutions and assigns a value between 0 and 1, where 0 indicates random labeling and 1 indicates identical cluster solutions (robust to a permutation of labels, adjusted for greater fit by chance at higher *k*). Note that, unlike previous applications (Yeo et al. 2011), we cross-validated AMI at the participant level rather than partitioning at the searchlight level.

Separate parcellations were obtained for each attention task condition to ensure the clustering algorithm did not attenuate task effects. The cluster analysis yielded qualitatively similar surface parcellations for data from both the behavior attention task and the taxonomy attention task, however the behavior attention task tended toward more reproducible solutions at higher *k*. Note that clustering cortical searchlights according to the pairwise neural distances between a certain set of experimental conditions should not be expected to yield a generally valid parcellation for the entire brain. Furthermore, although spatial smoothing, overlapping searchlights, and hyperalignment induce spatial correlations, there is nothing intrinsic to the clustering algorithm that ensures spatial contiguity (on the cortical surface) or bilaterality in the resulting parcellation.

The reproducibility analysis indicated local maxima at *k* = 2, 4, 14, 19, and 23 (Supplementary Fig. 4*A*), and these cluster solutions can then be mapped back to the cortical surface (Supplementary Figs. 4*B*, 5). All subsequent analyses were performed on regions of interest (ROIs) derived from the parcellation at *k* = 19 based on the behavior attention data. From these 19 areas tiling the entire cortical surface, 10 ROIs were selected comprising early visual areas, the ventral visual pathway, the dorsal visual pathway, and somatomotor cortex. These 10 ROIs corresponded to the areas of the brain with the highest inter-participant correlation of RDMs for both tasks (Supplementary Fig. 1*D*). Both the clustering algorithm and the reproducibility analysis are agnostic to any particular representational geometry or task effect (Kriegeskorte et al. 2009). ROIs were large, including on average 1,980 voxels (SD = 1,018 voxels; see Supplementary Table 2 for individual ROI extents).

### Correlations with target RDMs

For each ROI, we used the stimulus-evoked patterns of activation across all referenced voxels to compute neural RDMs for both attention conditions. We tested for task differences in Spearman correlation between the observed neural RDM and the target RDMs. To test this, we first constructed a linear mixed-effects model to predict Spearman correlations with the categorical target RDMs using Task, Target RDM, and ROI, and their two- and three-way interactions as fixed effects, with Participant modeled as a random effect (random intercepts). The Task variable captured the two attentional task conditions, Target RDM represented the behavioral and taxonomic category target RDMs, and ROI represented the 10 regions of interest. Mixed-effects modeling was performed in R using *lme4* (Bates et al. 2015). Statistical significance was assessed using a Type III analysis of deviance.

To assess the statistical significance of differences in Spearman correlation as a function of attention task for each ROI, nonparametric randomization tests were performed in which the mean difference in Spearman correlation (the test statistic) was computed for all possible permutations of the within-participants attention task assignments (2^12^ = 4,096 permutations, two-sided exact test). This approach permutes the signs of the within-participant task differences across participants. This tests the null hypothesis that there is no reliable effect of the attentional task manipulation across participants (task assignment is exchangeable within participants under the null hypothesis), and approximates a paired *t*-test where participant is modeled as a random effect. This nonparametric significance test was used for all subsequent tests of the attentional manipulation within ROIs. For visualization, mean Spearman correlations are plotted with bootstrapped 95% confidence intervals. Bootstrapping was performed at the participant level; that is, confidence intervals were constructed by sampling (with replacement) from the within-participant task differences in Spearman correlation to respect the within-participants comparison (Loftus and Masson, 1994). Note that Spearman correlation accommodates ties in a way that can be problematic when comparing RDMs with numerous ties (e.g., categorical RDMs, like those used here), particularly relative to RDMs with more continuous dissimilarity values (i.e., fewer ties; Nili et al., 2014). However, this does not negatively impact the present analysis where we compute Spearman correlations using the same RDM in different task contexts. To more directly interface with the searchlight analysis, we used the same standardized rank regression to examine task-related differences in representational geometry. This approach models the neural representational geometry of an ROI as a weighted sum of the behavioral category and taxonomic category target RDMs.

To ensure that our findings were not biased by the unsupervised functional parcellation method used to identify cortical ROIs with consistent representational geometries, we reproduced the above analysis in anatomically-defined ROIs. We extracted rough analogues of four key ROIs using the FreeSurfer cortical surface parcellation (Destrieux et al. 2010). The VT ROI was defined bilaterally as the conjunction of the fusiform gyrus (lateral occipitotemporal gyrus), collateral sulcus (medial occipitotemporal sulcus) and lingual sulcus, and lateral occipitotemporal sulcus parcels. IPS comprised the bilateral intraparietal sulci, transverse parietal sulci, and superior parietal lobules. PCS was defined bilaterally to include the postcentral gyrus, postcentral sulcus, and supramarginal gyrus, extending superiorly to *z* = 50 on the inflated cortical surface of the reference participant (in the hyperalignment algorithm) in MNI space. The vPC/PM ROI comprised the bilateral precentral gyri, central sulci, and subcentral gyri and sulci, similarly extending superiorly to *z* = 50. This superior boundary is roughly coterminous with the extension of the upper bank of the intraparietal sulcus anteriorly and was imposed on the PCS and vPC/PM ROIs to better match the functionally-defined ROIs reported above. Note hyperalignment effectively projects all participants’ data into the reference participant’s anatomical space.

Recent work (Walther et al. 2015) suggests that neural RDMs may be more reliably estimated by computing pairwise distances between conditions in a cross-validated fashion (i.e., across scanner runs). We re-computed the above analyses using alternative distance metrics and cross-validation schemes. We first estimated neural RDMs using Euclidean distance rather than correlation distance. Different distance metrics have different theoretical interpretations, and each has strengths and weaknesses. For example, correlation distance is susceptible to baseline shifts and noise, while Euclidean distance (and related metrics, such as Mahalanobis distance) is sensitive to regional-average differences in activation magnitude (Kriegeskorte et al. 2008; Walther et al. 2015). Next, we computed neural RDMs using leave-one-run-out cross-validation. Response patterns were estimated for each scanning run. Responses for four of the five runs for each attention task were averaged and pairwise distances were computed between each condition in the averaged runs and the left-out fifth run. This results in a neural RDM with nonzero distances in the diagonal cells. We then compared these neural RDMs with the categorical target RDMs using Spearman correlation, as described above. These analyses were performed post-hoc, in an exploratory fashion.

### Evaluating model fit

As evidenced by the searchlight analysis (Fig. 2), the target RDMs for taxonomy and observed behavior representation may differ in the extent to which they capture neural representational geometry. To address this, we evaluated differences in the fit of these models. However, although the target RDMs were sufficient to test our hypothesis, they cannot capture differences in the distances between behavioral and taxonomic categories; e.g., the animacy continuum (Connolly et al. 2012; Sha et al. 2015). To accommodate this type of geometry for behavior and taxonomy, we decomposed the categorical target RDMs into separate regressors for each between-category relationship. The number of regressors used was determined by the number of pairwise relationships between categories. The number of pairwise relationships between *n* categories is (*n* × (*n* – 1)) / 2. Therefore, for the four action categories, there are (4 × (4 – 1)) / 2 = 6 pairwise relationships. For the five animal categories, there are (5 × (5 – 1)) / 2 = 10 pairwise relationships. For example, the taxonomy model consists of a separate regressor for each within-category “box” along the diagonal of the taxonomic category target RDM (Fig. 1).

**Figure 1.**
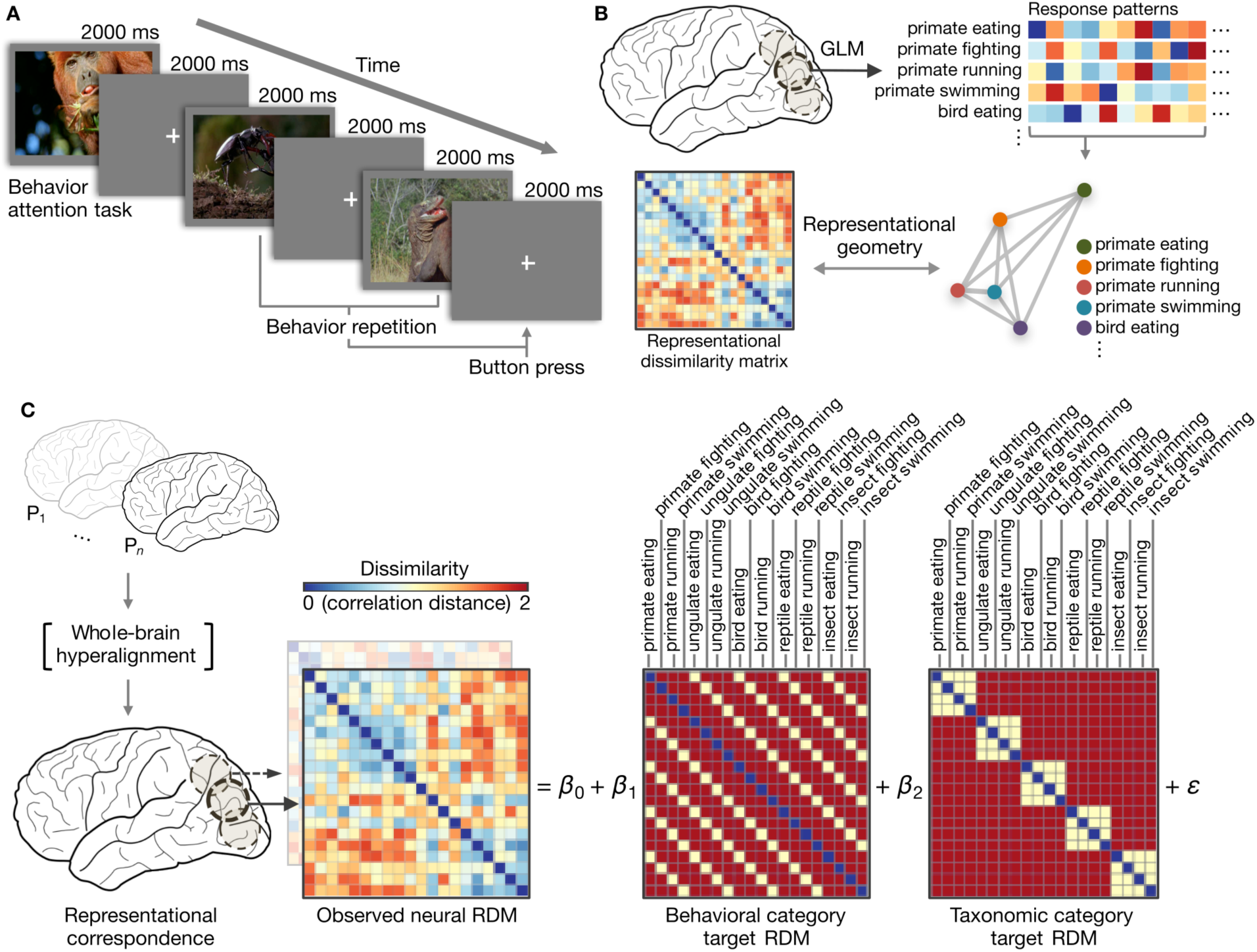
Experimental procedure and analytic approach. (*A*) Schematic of event-related design with naturalistic video clips of behaving animals (Supplementary Table 1, Supplementary Video 1). Participants performed a repetition detection task requiring them to attend to either animal taxonomy or behavior. (*B*) Stimulus-evoked response patterns for the 20 conditions were estimated using a conventional general linear model. The pairwise correlation distances between these response patterns describe the representational geometry (representational dissimilarity matrix; RDM) for a given brain area. (*C*) Whole-brain surface-based searchlight hyperalignment was used to rotate participants’ responses into functional alignment based on an independent scanning session (Supplementary Fig. 1). Following hyperalignment, the neural representational geometry in each searchlight was modeled as a weighted sum of models capturing the taxonomic and behavioral categories. Model RDMs were constructed by assigning correlation distances of 0 to identical conditions (the diagonal), correlation distances of 1 to within-category category distances, and correlation distances of 2 to between-category distances. Note that absolute distances assigned to these model RDMs are unimportant as only the ordinal relationships are preserved when using rank correlation metrics (e.g., Spearman correlation). Only the vectorized upper triangular of the RDMs (excluding the diagonal) are used. The observed neural representational geometry of a searchlight in posterolateral fusiform gyrus in a representative participant is used as an example. Supplementary Fig. 2 provides more detailed examples of searchlight representational geometries.

To evaluate these two flexible behavior and taxonomy models, in each ROI and each participant we computed the coefficient of partial determination (partial *R*^2^), then averaged these model fits over the two attention tasks (van den Berg et al. 2014). Partial *R*^2^ can be interpreted as the proportion of variance accounted for by one model controlling for any variance accounted for by the other model, and was computed separately for each attention task and then averaged across tasks within participants. We then computed the within-participants differences between the two models per ROI, and submitted these differences to a nonparametric randomization test to assess significance across participants. In the nonparametric randomization test, we permuted the direction of the within-participant difference in model fits across participants, testing the null hypothesis that there is no reliable difference in model fits across participants. The test statistic was the mean within-participants difference in model fit between the decomposed taxonomic and behavioral category models. Note, however, that partial *R*^2^ is biased toward more complex models (in this case, the taxonomy model), so we corroborated this analysis using the Akaike information criterion (AIC), which penalizes more complex models. We computed the difference in AIC for the six- and 10-regressor models for each attention task condition within each participant, then averaged across the attention tasks. These differences in AIC were assessed statistically using an exact test permuting the sign of the difference.

### Task-related differences in representational distances

Next, we probed for task-related differences in representational distances directly. Note however that certain pairwise distances (e.g., the distance between neural representations of a bird eating and an insect fighting) would not be hypothesized to change in a meaningful way as a function of our task manipulation (see, e.g., the diagonal distances in Fig. 4*B*). For this reason, we constrained our analysis to only within-category pairwise distances (cells of the RDM). Correlation distances were converted to Fisher-transformed correlations prior to statistical testing. Rather than averaging the pairwise distances across cells of the target RDM within each participant, cells corresponding to particular pairwise distances were included as a random effect (as per an items analysis; Baayen et al. 2008). We constructed a linear mixed-effects model to predict observed correlation distances based on Task, Category, and ROI, and their two- and three-way interactions as fixed effects, with Participant and Cell as random effects (random intercepts). Task represented the attentional task condition, Category represented the category relationship (within-behavior or within-taxon), ROI indicated the 10 ROIs reported above, and Cell indicated particular cells (pairwise relationships) of the target RDM. Statistical significance was assessed using a Type III analysis of deviance. Nonparametric randomization tests were used to assess task-related differences in mean within-category correlation distances within each ROI.

### Visualizing neural representational space

To visualize task-related changes in representational geometry, we used multidimensional scaling (Kriegeskorte et al. 2008b). For a given ROI, we first computed 40 × 40 neural RDMs based on the 20 conditions for both attention tasks and averaged these across participants. Note that between-task differences in the 40 × 40 neural RDM may be difficult to interpret, as the attentional task manipulation is confounded with scanner runs (Henriksson et al., 2015). However, we would not expect simple run differences to result in the observed attentional differences in representational distances for both behavior and taxonomy. To visualize task-related changes in observed action representation, we computed an 8 × 8 distance matrix comprising the mean between-behavior distances within each taxonomic category (as in Fig. 4). For taxonomy representation, we computed the average between-taxon distances within each behavioral category to construct a 10 × 10 matrix. Distances were computed between conditions for both tasks (e.g., resulting in a single 8 × 8 distance matrix rather than two separate 4 × 4 matrices for behavior representation) to ensure that distances for both attention tasks were on the same scale.

These distance matrices were then submitted to metric multidimensional scaling implemented in scikit-learn (Pedregosa et al. 2011). In the case of behavior representation, for example, this resulted in eight positions in a two-dimensional space. However, because we were interested in the overall task-related expansion between conditions (and less concerned with, e.g., the distance between one condition in one attention task and another condition in the other attention task), the positions in the resulting two-dimensional solution were then split according to attention task, and the Procrustes transformation (without scaling) was used to best align the conditions within one attention task to another. This transformation preserves the relationships between conditions within each task and captures the overall attentional expansion of between-category distances.

## Results

### Behavioral performance

Participants were highly accurate in detecting the sparse repetition events for both attention conditions (mean accuracy for animal attention condition = 0.993, SD = 0.005; mean accuracy for behavior attention condition = 0.994, SD = 0.005). There was no significant task-related difference in either accuracy (*t*(11) = 0.469, *p* = 0.648), or signal detection theoretic measures of sensitivity (*t*(11) = 0.116, *p* = 0.910) and bias (*t*(11) = 0.449, *p* = 0.662) adjusted for logistic distributions (according to Kane et al. 2007, p. 617). Response latencies for repetition trials where participants responded correctly did not significantly differ between the behavior attention and taxonomy attention tasks (paired *t*-test: *t*(11) = 0.015). However, the scanner protocol was not designed to robustly measure response times, as there were only four repetition events per run and participants did not respond to non-repetitions.

### Searchlight analysis

We applied representational similarity analysis using surface-based searchlights to map areas of the brain encoding information about animal taxonomy and behavior. Neural representational dissimilarity matrices (RDMs) were computed based on the pairwise correlation distances between hyperaligned stimulus-evoked response patterns for the 20 conditions (Fig. 1*B*). We modeled the neural representational geometry as a weighted sum of two categorical target RDMs reflecting the experimental design: a behavioral category target RDM and a taxonomic category target RDM (Fig. 1*C*).

We first identified clusters of searchlights where the neural representational geometry reflected the categorical target RDMs for both attention conditions. Regression coefficients for the behavioral category target RDM were strongest in lateral occipitotemporal (LO) cortex, in the dorsal visual pathway subsuming posterior parietal, intraparietal sulcus (IPS), motor and premotor areas, and in ventral temporal (VT) cortex (Fig. 2*A*). Regression coefficients for the animal taxonomy target RDM were strongest in VT, LO, and posterior parietal cortices, as well as left inferior and dorsolateral frontal cortices.

**Figure 2.**
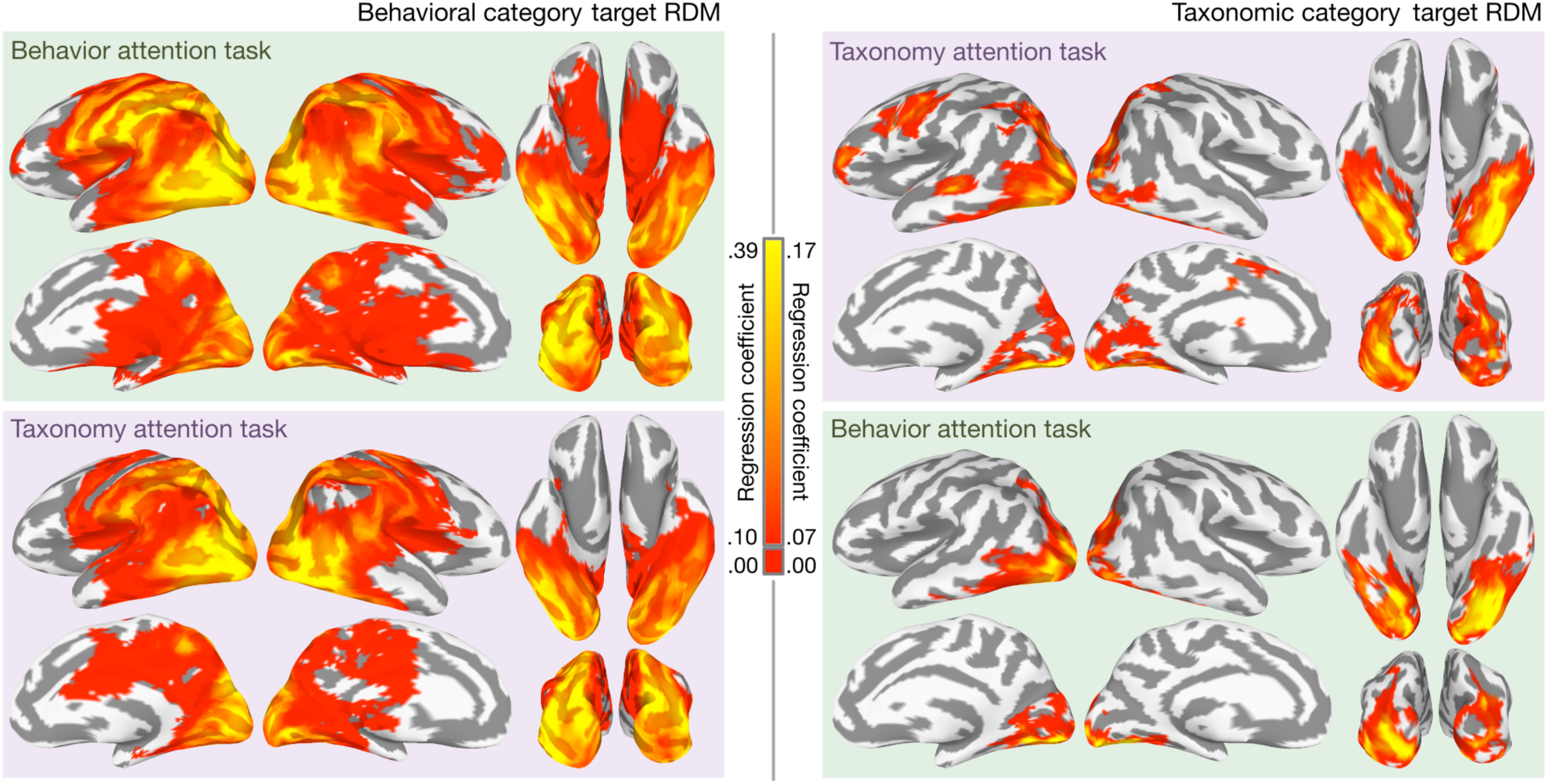
Mapping representations of animal behavior and taxonomy for both tasks. Significant searchlight regression coefficients for the behavioral category target RDM (left) and the taxonomic category target RDM (right) are mapped onto the cortical surface for both attention conditions. Cluster-level significance was assessed at the group level using TFCE and maps are thresholded at cluster-level *p* < .05 (nonparametric one-sided test, corrected for multiple comparisons). For searchlights surviving cluster-level significance testing, the mean regression coefficient across participants is plotted. All colored searchlights exceed the cluster-level threshold of statistical significance across participants, corrected for multiple comparisons using TFCE; searchlights not surviving cluster-level significance testing are not colored. Note that regression coefficients for behavior representation and taxonomy representation are plotted with different color scales to better visualize the distribution of coefficients. Regression coefficients less than 0.10 for the behavioral category target RDM and less than 0.07 for the taxonomic category target RDM are plotted as red. See Supplementary Fig. 2 for qualitatively similar searchlight classification maps, and Supplementary Fig. 3 for difference maps.

Based on previous work (e.g., Çukur et al. 2013), we hypothesized that attending to a particular type of semantic information would enhance task-relevant representational distinctions in searchlights encoding taxonomic and behavioral category information throughout the cortex. Globally, attending to behavior or taxonomy increased the regression coefficients for the target RDMs corresponding to the attended categories. Attending to behavior increased the number of searchlights with significant regression coefficients for the behavioral category target RDM from 11,408 to 14,803 (corrected for multiple comparisons). We next tested whether attentional allocation increased the mean regression coefficient for the behavioral category target RDM in searchlights containing information about the behavioral categories. When considering regression coefficients for the behavioral category target RDM in all searchlights surviving multiple comparisons correction for either attention task, attending to animal behavior significantly increased the mean regression coefficient from 0.100 to 0.129 (*p* = .007, nonparametric randomization test). Attending to taxonomy increased the number of searchlights with significant regression coefficients for the taxonomic category target RDM from 1,691 to 3,401. For searchlights surviving multiple comparisons correction for either task, regression coefficients for the taxonomic category RDM increased significantly from 0.049 to 0.071 (*p* = .017, nonparametric randomization test). A linear SVM searchlight classification analysis, in which we used leave-one-category-out data folding for cross-validation (Supplementary Fig. 2), resulted in qualitatively similar maps, suggesting the results presented in Fig. 2 are not driven solely by low-level visual properties of particular stimuli (although low-level visual properties may still covary with condition).

### Regions of interest

We hypothesized that task demands may alter representational geometry across larger cortical fields than captured by the relatively small searchlights. We tested our hypothesis in large ROIs defined by shared searchlight representational geometry. We applied an unsupervised clustering algorithm to the searchlight representational geometries to parcellate cortex into ROIs and used a relatively reproducible parcellation with 19 areas (Supplementary Fig. 4). We interrogated 10 ROIs with high inter-participant similarity of searchlight representational geometry subtending the dorsal and ventral visual pathways (Fig. 3*B*, Supplementary Fig. 1). The 10 ROIs were labeled as follows: posterior early visual cortex (pEV), inferior early visual cortex (iEV), superior early visual cortex (sEV), anterior early visual cortex (aEV), lateral occipitotemporal cortex (LO), ventral temporal cortex (VT), occipitoparietal and posterior parietal cortex (OP), intraparietal sulcus (IPS), left postcentral sulcus (left PCS), and ventral pericentral and premotor cortex (vPC/PM).

**Figure 3.**
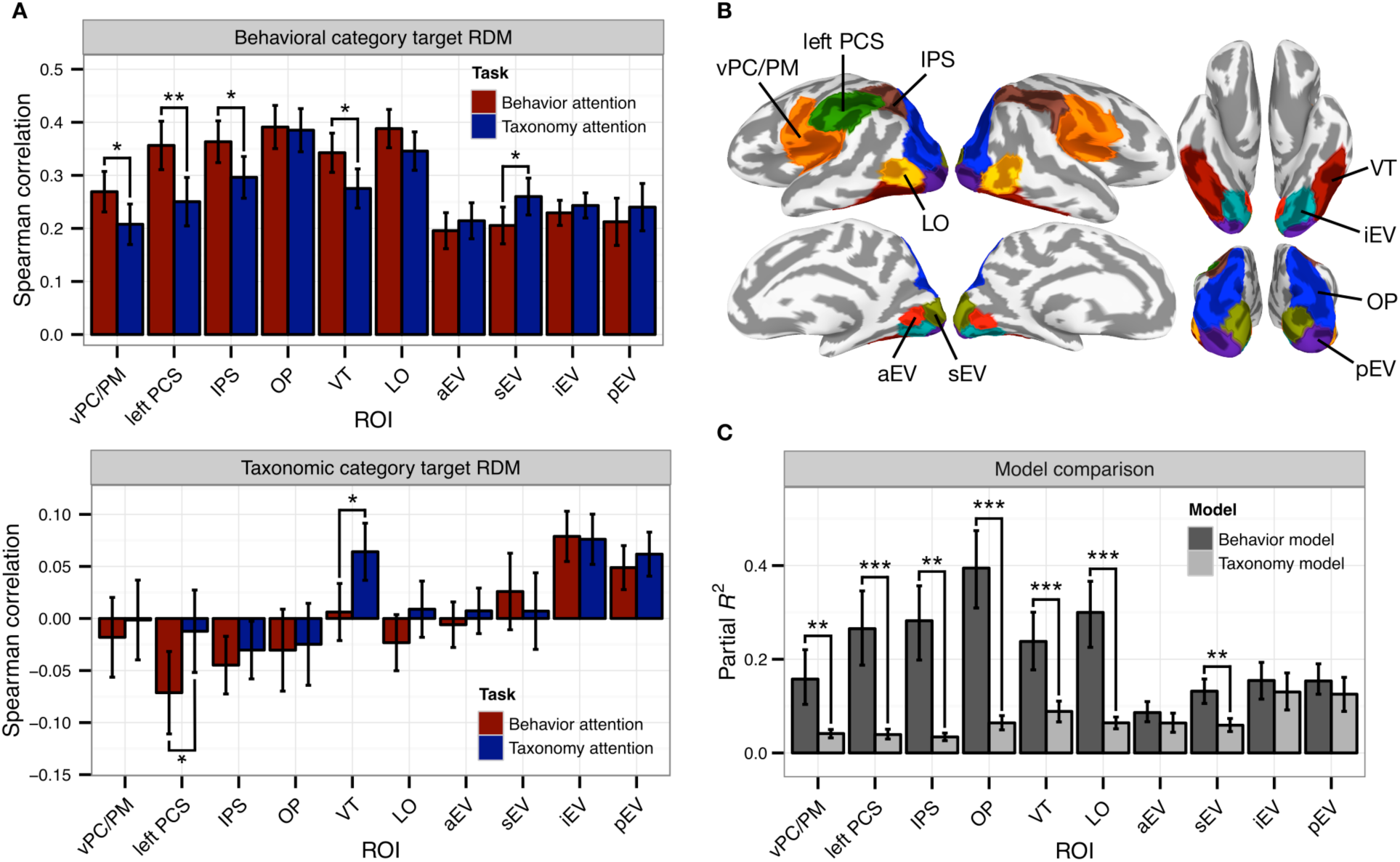
Attention alters representational geometry in functionally-defined ROIs. (*A*) Task differences in Spearman correlation between neural RDMs and the behavioral and taxonomic category target RDMs (see Supplementary Table 2 for results for all 19 clusters). Participants were bootstrap resampled to construct 95% confidence intervals for within-participant effects. Supplementary Fig. 7 presents key findings reproduced in anatomically-defined ROIs. Supplementary Fig. 8 depicts qualitatively similar results using standardized rank regression rather than Spearman correlation. See Supplementary Fig. 9 for similar analyses computed using alternative pairwise distance metrics and cross-validation schemes. (*B*) Ten functional ROIs identified by parcellating the cerebral cortex based on representational geometry. (*C*) Comparison of model fit for the six-regressor behavior model and 10-regressor taxonomy model. \**p* < .05, \*\**p* < .01, \*\*\**p* < .001, two-sided nonparametric randomization test.

For each ROI, we measured the Spearman correlation between the observed neural RDM and the two categorical target RDMs for each task, to test whether task demands altered neural representational geometry (Fig. 3*A*). A linear mixed-effects model yielded significant main effects for ROI (*χ*^2^(9) = 115.690, *p* < .001) and Target RDM (*χ*^2^(9) = 69.640, *p* < .001), and a significant Target RDM × ROI interaction (*χ*^2^(9) = 112.442, *p* < .001). The Task × ROI interaction was also significant (*χ*^2^(9) = 23.301, *p* = .006), suggesting that the task manipulation more strongly affected correlations in certain ROIs than others. Note however that differences due to the task manipulation across ROIs could be driven by different noise levels in different ROIs (Diedrichsen et al. 2011). Finally, the three-way Task × Target RDM × ROI interaction was significant (*χ*^2^(9) = 22.034, *p* = .009), motivating the following within-ROI tests. Nonparametric randomization tests revealed that attending to animal behavior increased correlations between the observed neural RDM and the behavioral category target RDM in vPC/PM (*p* = .026), left PCS (*p* = .005), IPS (*p* = .011), and VT (*p* = .020). A decrease in the categoricity of behavior representation was observed in sEV when participants attended to behavior (*p* = .032). Attending to animal taxonomy increased correlations between the observed neural RDM and the taxonomic category target RDM in VT (*p* = 0.010) and left PCS (*p* = .036). The effect in left PCS was driven by a negative correlation in the behavior attention task that was abolished when attention was directed at taxonomy. To ensure that these effects were not biased by the functional parcellation technique used to define functional ROIs, we reproduced key findings in anatomically-defined ROIs (Supplementary Fig. 7). To better interface with the searchlight results, we reproduced qualitatively similar findings using the standardized rank regression technique used in the searchlight analysis (Supplementary Fig. 8). Unlike computing Spearman correlation separately per RDM, this approach allocates variance in neural representational geometry to both RDMs. This analysis yielded generally greater regression coefficients for the taxonomic category RDM, suggesting that the low and negative correlations observed using Spearman correlation with the taxonomic category RDM (e.g., in left PCS) may be due to variance in representational geometry related to the behavioral categories. Attending to behavior significantly enhanced behavioral category representation in anatomically-defined vPC/PM, bilateral PCS, and VT ROIs, while attending to taxonomy strongly enhanced taxonomic category representation in VT and weakly in vPC/PM. We also reproduced qualitatively similar findings using Euclidean distance rather than correlation distance and using leave-one-run-out cross-validation in constructing the neural RDMs (Supplementary Fig. 9). Supplementary Tables 2 and 3 present task differences in Spearman correlation for all 19 parcels returned by the cluster analysis and all anatomically discontiguous parcels, respectively.

Unexpectedly, behavioral category representation was found to be considerably stronger and more prevalent than taxonomic category representation. To test this formally, we next evaluated how well full representational models of animal taxonomy and behavior fit the neural representational geometry in each ROI. The model RDMs used above tested our experimental hypothesis but do not capture the geometry of distances between behavioral or taxonomic categories; e.g., the animacy continuum (Connolly et al. 2012; Sha et al. 2015). To accommodate this type of geometry for behavior and taxonomy, we decomposed the categorical target RDMs into separate regressors for each pairwise between-category similarity (six regressors for behavior model, 10 for the taxonomy model). To evaluate these two flexible behavior and taxonomy models, in each ROI we estimated the coefficient of partial determination (partial *R*^2^) and AIC separately for each model and attention task within each participant, then averaged these model fits over the two attention tasks. The six-regressor behavior model captured on average over 2 times more variance (adjusted *R*^2^) than the single-regressor behavioral category target RDM in LO, VT, OP, IPS, left PCS, and vPC/PM, suggesting that some behaviors are more similar to each other than are others. The 10-regressor taxonomy model accounted for well over 4 times more variance than the single-regressor taxonomic category target RDM in pEV, iEV, and VT. Based on nonparametric randomization tests, partial *R*^2^ for the behavior model significantly exceeded that of the animal taxonomy model in sEV, LO, VT, OP, IPS, left PCS, and vPC/PM (Fig. 3*C*), and AIC for the behavior model was significantly lower for all 10 ROIs. Surprisingly, the behavior model accounted for over 2.5 times more variance in VT neural representational geometry than did the taxonomy model (behavior model: 23.8% of variance; taxonomy model: 8.8% of variance).

Although the initial ROI analysis demonstrated that the attention task alters overall neural representational geometry to more closely resemble the categorical target RDMs, it does not directly quantify changes in representational distances. To test task-related changes in representational distances more explicitly, we isolated cells of the neural RDM capturing distances between two conditions that differed on one dimension and were matched on the other; i.e., different behaviors performed by animals from the same taxonomic category, or animals of different taxonomic categories performing the same behavior (Fig. 4*A*). Although we hypothesized that attention enhances task-relevant representational distinctions as depicted in Fig. 4*B* (40, 41), note that diagonal distances do not change; that is, the effect of attention on distances between conditions that differ on both dimensions is ambiguous. Thus, focusing on the correlation distances between pairs of conditions that differ on only one dimension affords a less confounded examination of the effects of attention. A significant increase in, e.g., between-taxon correlation distances within each behavior (Fig. 4*A*, red) when attending to behavior can also be interpreted as a decrease in within-taxon distances when attending to taxonomy; therefore, we refer to this effect as enhancing task-relevant representational distinctions. A linear mixed-effects model yielded significant main effects for ROI (*χ*^2^(9) = 66.850, *p* < .001) and Category (within-behavior or within-taxon category relationship; *χ*^2^(9) = 13.047, *p* < .001), as well as a significant ROI × Category interaction (*χ*^2^(9) = 165.725, *p* < .001). Most importantly, this analysis revealed a significant three-way Task × Category × ROI interaction (*χ*^2^(9) = 33.322, *p* < .001), motivating the following within-ROI tests. Nonparametric randomization tests indicated that attention significantly enhanced task-relevant representational distinctions for both groups of distances in left PCS (between-taxon, within-behavior distances: *p* = .002; between-behavior, within-taxon distances: *p* = .010) and VT (between-taxon, within-behavior distances: *p* = .028; between-behavior, within-taxon distances: *p* = .009). Attention significantly enhanced task-relevant between-taxon distances within behaviors in vPC/PM (*p* = .007), effectively collapsing taxonomic distinctions when attending to behavior. An inverted task effect was observed in sEV (between-taxon, within-behavior distances: *p* = .028). Supplementary Tables 4 and 5 present the task enhancement of representational distances for all 19 parcels returned by cluster analysis and all anatomically discontiguous parcels, respectively. The expansion of distances between attended category representations is illustrated with multidimensional scaling of the representational geometries in left PCS and VT (Fig. 4*C*).

**Figure 4.**
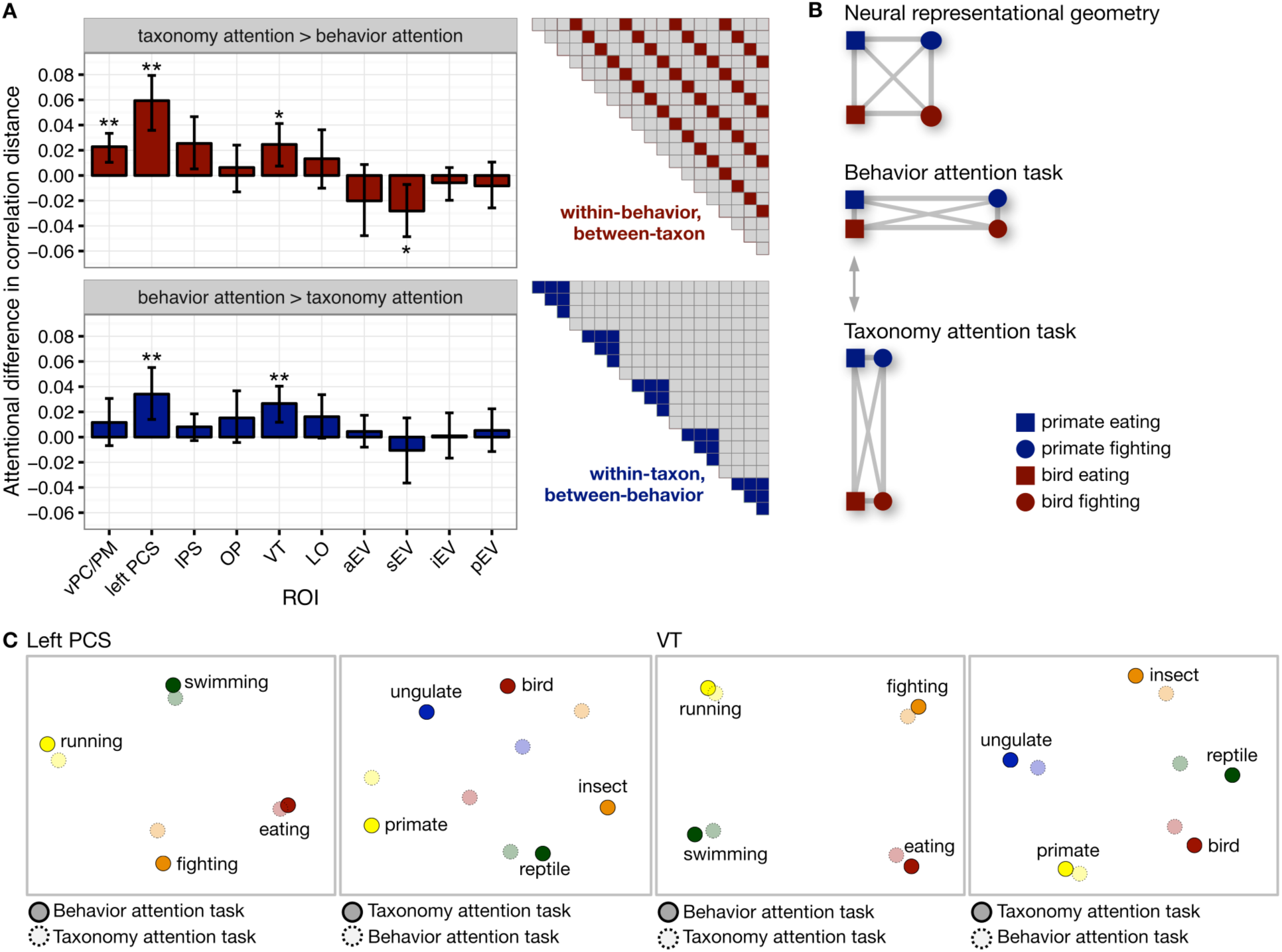
Attention enhances the categoricity of neural responses patterns. (*A*) Enhancement of within-category distances for both behavioral and taxonomic categories based on the attention task (see Supplementary Table 3 for results for all 19 clusters). Error bars indicate bootstrapped 95% confidence intervals for within-participants task differences (bootstrapped at the participant level). (*B*) Schematic illustrating how neural distances are expanded along the behaviorally relevant dimensions while task-irrelevant distances are collapsed (Nosofsky 1986; Kruschke 1992). (*C*) Multidimensional scaling (MDS) solutions for left PCS and VT depict the attentional expansion of between-category distances. \**p* < .05, \*\**p* < .01, two-sided nonparametric randomization test.

## Discussion

The present study was motivated by the following question: How does attention prioritize certain semantic features of a complex stimulus in service of behavioral goals? We hypothesized that attention may enhance certain features of semantic information encoded in distributed neural populations by transiently altering representational geometry (Kriegeskorte and Kievit 2013). Our findings provide neural evidence for psychological theories of attentional deployment in categorization (Shepard, 1964; Tversky, 1977; Nosofsky 1986; Kruschke 1992) by demonstrating that attention selectively increases distances between stimulus-evoked neural representations along behaviorally relevant dimensions. To expand on prior work examining early visual (e.g., orientation, contrast, color, motion direction; Serences and Boynton 2007; Jehee et al. 2011; Brouwer and Heeger 2013; Sprague and Serences 2013) and object category (Peelen et al. 2009; Çukur et al. 2013; Harel et al. 2014; Erez and Duncan 2015) representation, we used dynamic, naturalistic stimuli to demonstrate that attention alters the representation of both animal taxonomy and behavior according to a similar principle.

The neural representation of animal taxonomy and behavior changed significantly with task. When participants attended explicitly to animal behavior, the categoricity of observed action representation increased most dramatically in premotor, pericentral, and postcentral somatomotor areas supporting action and goal recognition (Oosterhof et al. 2010, 2012, 2013; Rizzolatti and Sinigaglia 2010), intraparietal areas implicated in executive control (Petersen and Posner 2012), and VT. The left-lateralization of this effect is consistent with generally left-lateralized representation of action concepts in the brain (Noppeney 2008; Watson et al. 2013). In the current study, we cannot rule out the possibility that attending to behavior enhances the representation of low-level motion-related features of the stimulus more so than higher-level semantic representations. However, we note that retinotopic visual areas driven primarily by motion energy (Nishimoto et al. 2011; Huth et al. 2012) and early areas exhibiting robust representation of animal behavior (e.g., LO and OP) were not strongly modulated by the task manipulation. Attending to animal taxonomy increased the categoricity of animal representation in VT, consistent with accounts of neural representation of animals and objects (Connolly et al. 2012; Grill-Spector and Weiner 2014; Sha et al. 2015), as well as left PCS, but not in lateral occipitotemporal or early visual areas. Note that attending to behavior induced a negative correlation for the taxonomic category target RDM in left PCS, while attending to taxonomy abolished this effect. This negative correlation when attending to behavior could be driven by increased distances between behavior representations within each animal taxon. Behavior and taxonomy representation observed in unexpected regions such as anterior prefrontal cortex using the searchlight approach may be due to both the richness of the information conveyed by naturalistic stimuli and the categorization and working-memory components of the task. The relative magnitudes of task-related and stimulus-driven contributions to representational geometry varied across cortical areas. Overall, attending to animal behavior, as compared to when participants attended to animal taxonomy, increased correlations with the behavioral category RDM from .25 to .33 on average in the three frontoparietal ROIs, with increases in correlation ranging from .06–.11 or 23–42%. Significant correlation between the taxonomic category RDM and the VT neural RDMs was observed only when participants attended to animal taxonomy.

Performing a categorization task requiring attention to either animal taxonomy or behavior enhances the categoricity of neural representations by accentuating task-relevant representational distinctions. Our results demonstrate that attentional allocation sculpts representational geometry in late-stage sensorimotor areas; this effect was not observed in early perceptual areas. This is in line with electrophysiological work in macaques demonstrating that object categorization training increases the precision of response selectivity for task-relevant stimulus features in cortical areas thought to support perceptual processing (i.e., temporal cortex; Sigala and Logothetis 2002). In a related series of reports, Peelen and colleagues (2009, 2011, 2014) have suggested that visual search (for objects such as humans and vehicles) is facilitated by the activation of task-relevant representational templates in late-stage visual areas. Our findings support this hypothesis in the context of a finer-grained taxonomic categorization task and suggest that this framework may extend beyond object detection to more abstract representational templates of observed actions. More generally, our results demonstrate that the representational geometry of semantic information in systems such as VT and somatomotor cortex is dynamic and actively tuned to behavioral goals, rather than being solely a reflection of static conceptual knowledge.

Behavioral performance was effectively at ceiling for both attention tasks, suggesting that although participants were compliant, the task may not have elicited strong attentional deployment. Furthermore, to reduce the impact of behavioral responses on the MRI data, participants were not required to respond to non-repetitions, and therefore submitted very few (e.g., four) behavioral responses per run. These limitations prevent us from making claims relating the magnitude of attentional deployment to the size of changes in representational geometry and examining trial-by-trial relationships between behavior and representational geometry. A more demanding attentional task may further enhance changes in representational geometry and reveal a more extensive cortical system that is modulated by attention. Parametric variation of attentional demand may allow quantification of the effect of attention on representational geometry. The magnitude of attentional deployment in naturalistic paradigms with more complex goals is difficult to vary systematically and may be relatively low compared to psychophysical paradigms employing controlled stimuli. Furthermore, we did not include a “baseline” or “no attention” task condition in the present study, as it is not clear what would constitute an appropriate “baseline” task in natural vision paradigms given the difficulty of controlling spontaneous allocation of attention to meaningful, dynamic stimuli. Because of these considerations, the significance of our findings rests on the relative differences between the behavior and taxonomy attention tasks.

Numerous visual areas coded for both taxonomy and behavior, suggesting these two types of information are encoded in distributed population codes in a superimposed or multiplexed fashion (Grill-Spector and Weiner 2014; Haxby et al. 2014). However, the behavior model accounted for notably more variance in neural representation throughout the cortex than the taxonomy model—even in areas typically associated with animal category representation, such as VT (Connolly et al. 2012; Sha et al. 2015). The dominance of behavior in the representational geometry of behaving animals may be related to the prevalence of biological motion energy information when viewing naturalistic video stimuli (Huth et al. 2012; Russ and Leopold 2015). Work by others shows that lateral fusiform cortex responds strongly to dynamic stimuli that depict agentic behavior with no biological form (Grossman and Blake 2002; Gobbini et al. 2007), and biological motion and social behaviors drive responses in face-selective temporal areas in the macaque (Russ and Leopold 2015). Future work can use eye-tracking and neurally-inspired motion-energy models (e.g., Nishimoto et al. 2011) to examine how viewing time, gaze patterns, and motion information contribute to observed action representation and how low-level stimulus properties, such as simple and biological motion energy, interact with endogenous attention.

By design, there was considerable heterogeneity both in the exemplar animals within each taxonomic category and the exemplar actions within each behavioral category. For example, the primate category included different species, with stimuli depicting a chimpanzee eating a fruit and a macaque swimming in a hot spring. However, behavioral categories were similarly heterogeneous, grouping, for example, the bonobo eating a fruit with stimuli depicting a hummingbird feeding from a flower and a caterpillar eating its own eggshell. The visual heterogeneity of the category exemplars attests to the top-down category structure imposed on the stimuli by the task demands. There is some behavioral evidence that actions (or verbs), similarly to objects (typically nouns; Rosch 1978), adhere to a hierarchical category structure with a “basic” (or most frequent, prototypical) intermediate level (Abbott et al. 1985; Rifkin 1985; Fellbaum 1990; Morris and Murphy 1990). The behavioral and taxonomic categories used here are at an intermediate level but may not be at a putative basic level of the semantic category hierarchy. Verb hierarchies, however, are qualitatively different from noun hierarchies, with a “more shallow, bushy structure” and fewer hierarchical levels (Fellbaum 1990), making it difficult to match the level of taxonomic and behavioral categories across their respective semantic hierarchies. Moreover, it is unclear to what extent neural representation (as accessible using fMRI) reflects the primacy of basic-level categories reported behaviorally (cf. Connolly et al. 2012).

The neural representation of observed animal behaviors (and observed actions more generally) may differ qualitatively from the neural representation of animal taxonomy. The stronger correlation of neural representational geometry with models of behavioral categories may be due to stronger neural responses to biological motion energy than to biological form (Huth et al. 2012; Russ and Leopold 2015). Furthermore, whereas taxonomic category can be ascertained quickly and does not change with time, observed behaviors evolve over time. The semantic content conveyed by behavior also differs considerably from that conveyed by taxonomy. For example, observed actions convey motor goals (Oosterhof et al. 2013; Rizzolatti and Sinigaglia 2010) and vary considerably in affective content. The rich, multidimensional information conveyed by dynamic stimuli depicting behaving animals in their natural environments may evoke responses in a variety of neural systems. Along these lines, the representation of taxonomy is also driven by semantic features such as animacy (Connolly et al. 2012; Sha et al. 2015) and perceived threat (Connolly et al. 2016). The neural representation of these features may rely on systems supporting affective and social cognition (Saxe 2006; Connolly et al. 2016).

The present study expands on work by (Brouwer and Heeger 2013) demonstrating that the neural color space in early visual areas becomes more categorical when participants perform a color naming task. Here, we use rich, naturalistic stimuli to demonstrate that task demands affect neural representations of animal taxonomy and behavior in a similar fashion in perceptual and somatomotor areas. The current findings also complement a recent study by Çukur and colleagues (2013) demonstrating that attending to a particular object category (humans or vehicles) shifts the semantic tuning of widely distributed cortical voxels toward that category, even when exemplars of that category are not present in the stimulus. Although the tuning shifts observed by Çukur and colleagues (2013) are consistent with a selective expansion of representational space, they may not be the exclusive underlying mechanism. For example, increased response gain, sharper tuning (Brouwer and Heeger 2013), and changes in the correlation structure among voxels (Chen et al. 2006; Miyawaki et al. 2008) may also contribute to the task-related differences we observe in distributed representation. Further work is needed to investigate the relative roles played by each of these candidate mechanisms in task-related changes of representational geometry measured from distributed response patterns. Nonetheless, our findings provide a direct demonstration of the task-related expansion of representational space hypothesized by Çukur and colleagues (2013) and extend the domain of attentional modulation from object categories to observed actions.

Scaling up the effects of attention on single neurons to population responses and multivoxel patterns of activity is an outstanding challenge. Top-down signals (Desimone and Duncan 1995; Baldauf and Desimone 2014) may bias how information is encoded by single neurons (Treue and Martinez Trujillo 1999; Sigala and Logothetis 2002) and at the population level by altering neuronal gain, tuning, and interneuronal correlations (Averbeck et al. 2006; Cohen and Maunsell 2009; Ruff and Cohen 2014; Downer et al. 2015) in order to optimize representational discriminability for downstream read-out systems. Our findings suggest a model whereby attention alters population encoding in late-stage perception so as to enhance the discriminability of task-relevant representational content. At an algorithmic level (Marr 1982), attention may tune a feature space of arbitrary dimensionality by dynamically altering population encoding. This mechanism could enhance behavioral performance by temporarily disentangling (DiCarlo et al. 2012) task-relevant representations and collapsing task-irrelevant content.

## Acknowledgments

We thank Kelsey Wheeler for help in collecting the video stimuli, Courtney Rogers for administrative support, and Nikolaus Kriegeskorte for providing many helpful comments in an open review.

### Author Contributions

S.A.N. and J.V.H. designed research; S.A.N. performed research; A.C.C., N.N.O., Y.O.H., J.S.G., M.V.D.O.C., J.G., and M.I.G. contributed analytic tools; S.A.N. analyzed data; and S.A.N. and J.V.H. wrote the paper.

## Supplementary Material

**Supplementary Figure 1.**
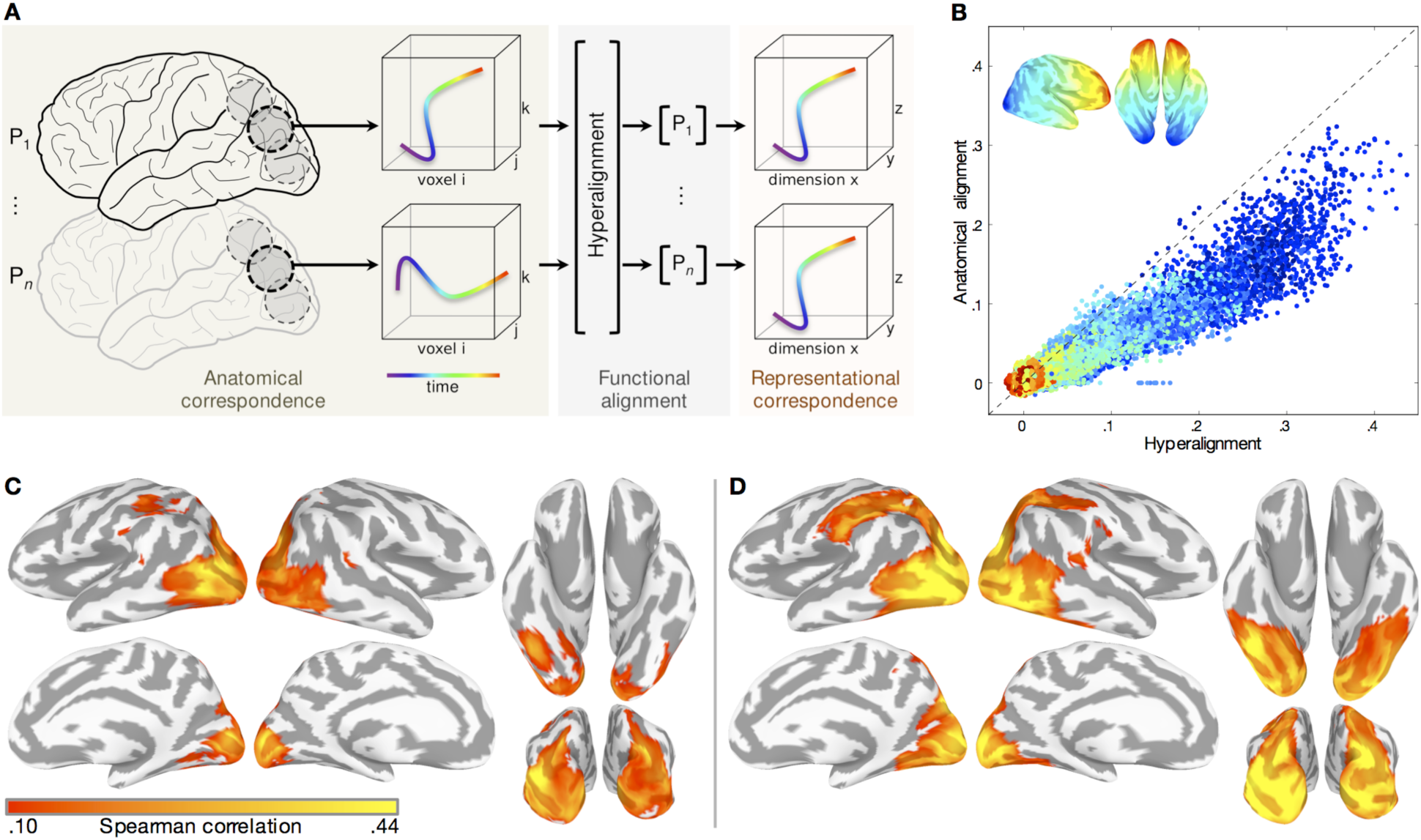
Whole-brain searchlight hyperalignment enhances representational correspondence across participants. (*A*) For each surface-based searchlight, the Procrustes transformation is used to rotate each participant’s time series of functional response patterns to the *Life* movie stimulus into a common space that maximizes representational correspondence across participants. These patterns are depicted as a trajectory of responses in a three-voxel space over time. (*B*) Each point in the scatterplot represents the average inter-participant Spearman correlation of RDMs for both attention tasks in a single searchlight. For each surface-based searchlight, the upper triangulars of the observed neural RDMs for both attention tasks were concatenated and pairwise Spearman correlations were computed between all participants. The vertical axis indicates Spearman correlation based on surface-based spherical alignment; the horizontal axis indicates Spearman correlation after surface-based searchlight whole-brain hyperalignment. Deviance from the identity line indicates a strong effect of alignment method on inter-participant similarity of RDMs. Searchlights are colored according their location on the posterior–anterior axis of the inflated cortical surface. (*C*) Inter-participant Spearman correlation of searchlight RDMs for both attention tasks using anatomical alignment thresholded at .10. (*D*) Average inter-participant Spearman correlation of searchlight RDMs after hyperalignment at the same threshold. Prior to hyperalignment, the maximum mean Spearman correlation was .32 in a searchlight superior to the left lateral occipital sulcus. Following hyperalignment, the maximum mean Spearman correlation was .44 in a searchlight in the left lateral occipital sulcus.

**Supplementary Figure 2.**
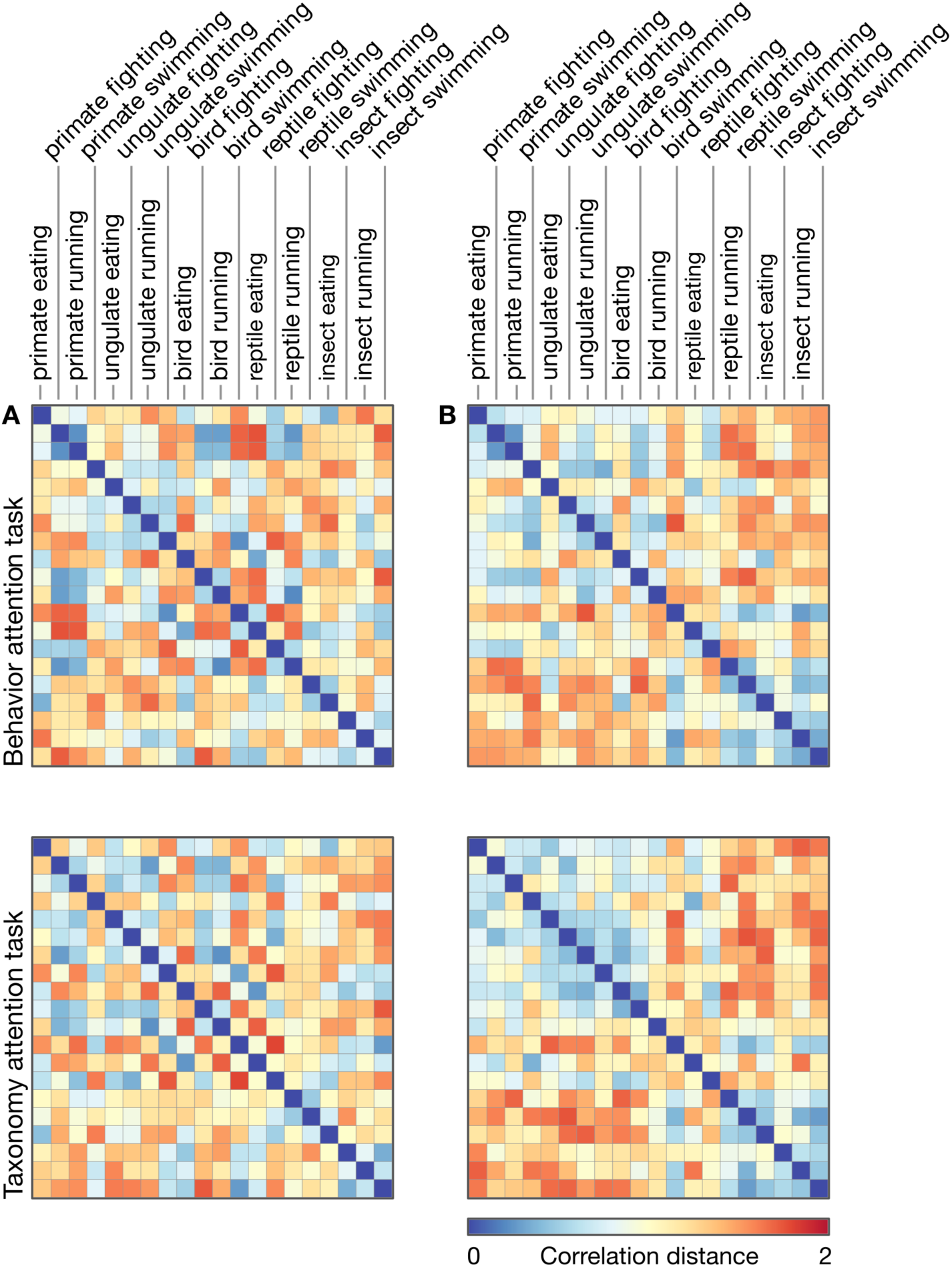
Example searchlight representational geometries. The observed neural RDMs for two example searchlights from a representative participant are plotted for the behavior attention task and the taxonomy attention task. (*A*) Observed neural RDM for a searchlight in left lateral occipitotemporal cortex (posterior middle temporal gyrus). Differences in behavioral category representation are reflected in repeated off-diagonal strips. (*B*) Observed neural RDM for a searchlight in left ventral temporal cortex (posterolateral fusiform gyrus). Taxonomic categories are represented according to an animacy continuum varying systematically in similarity from primates (most animate) to insects (least animate; Connolly et al. 2012; Sha et al. 2015). Example searchlights were not selected to convey an effect of the task manipulation.

**Supplementary Figure 3.**
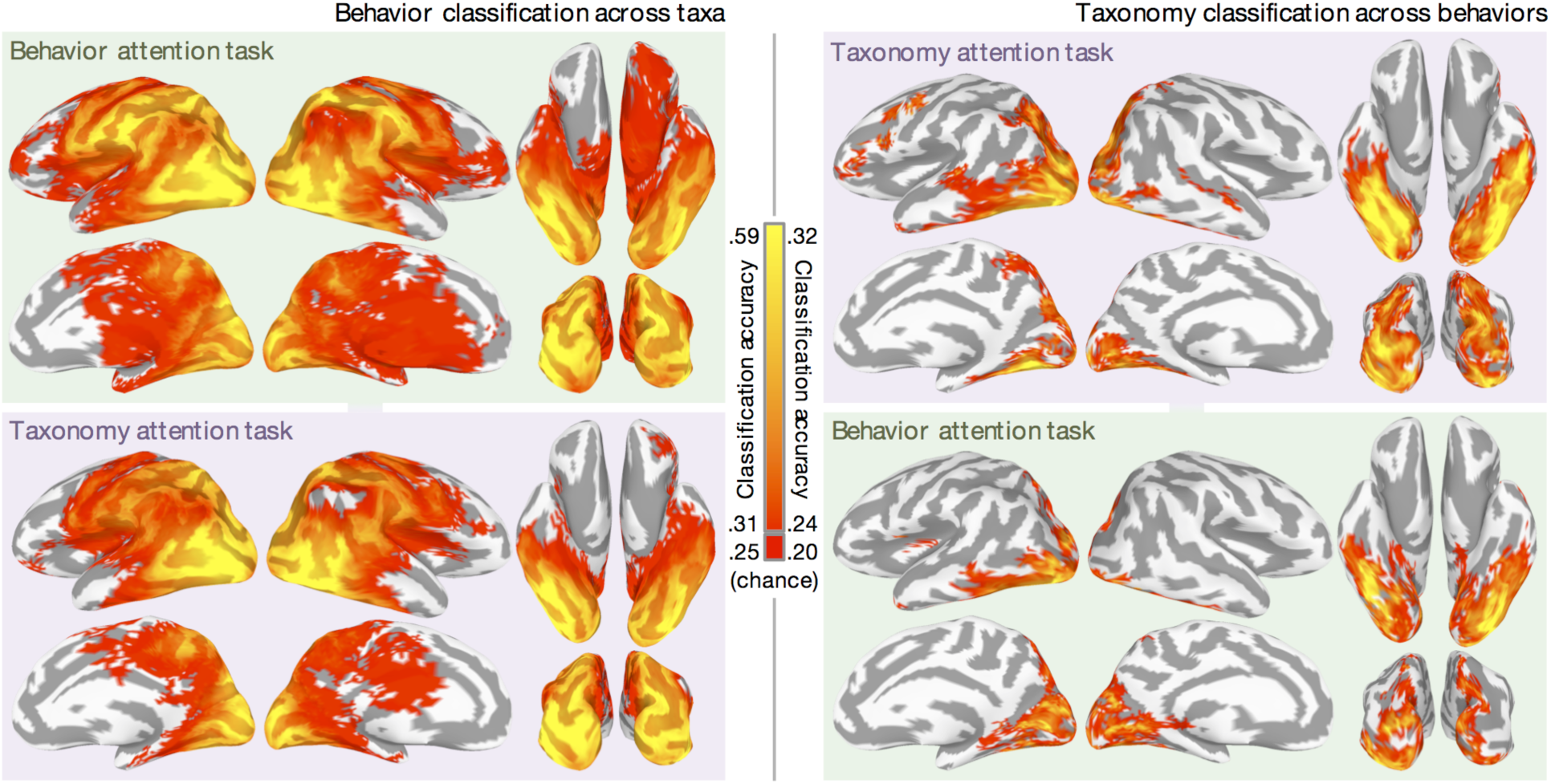
Effect of attention on searchlight classification of behavior and taxonomy. Cross-validation was implemented in the following leave-one-category-out fashion: classifiers discriminating the four behaviors (left) were trained on four of the five taxa, and tested on the left-out taxon; classifiers discriminating the five animal taxa (right) were trained on three of the four behaviors and tested on the left-out behavior. This procedure ensured that any information about animal behavior generalizes across animal taxa, and vice versa. Furthermore, classifiers in this cross-validation scheme are always tested on exemplar clips not in the training set, ensuring that classification accuracy is not based solely on low-level visual properties idiosyncratic to particular stimuli. Prior to classification, the GLM was computed separately for each run, yielding 20 beta parameters per run. The maps are qualitatively similar to the representational similarity regression maps reported in Fig. 2, with an average correlation of .83 across conditions prior to thresholding. Chance accuracy for four-class behavior classification is .25 and chance accuracy for five-class taxonomy classification is .20. Accuracies less than 0.31 for behavior classification and less than .24 for taxonomy classification are plotted as red. Maps are thresholded at *p* < .05 using TFCE, based on a null distribution of searchlight maps generated by permuting the labels of interest within each run and within each category of the crossed factor.

**Supplementary Figure 4.**
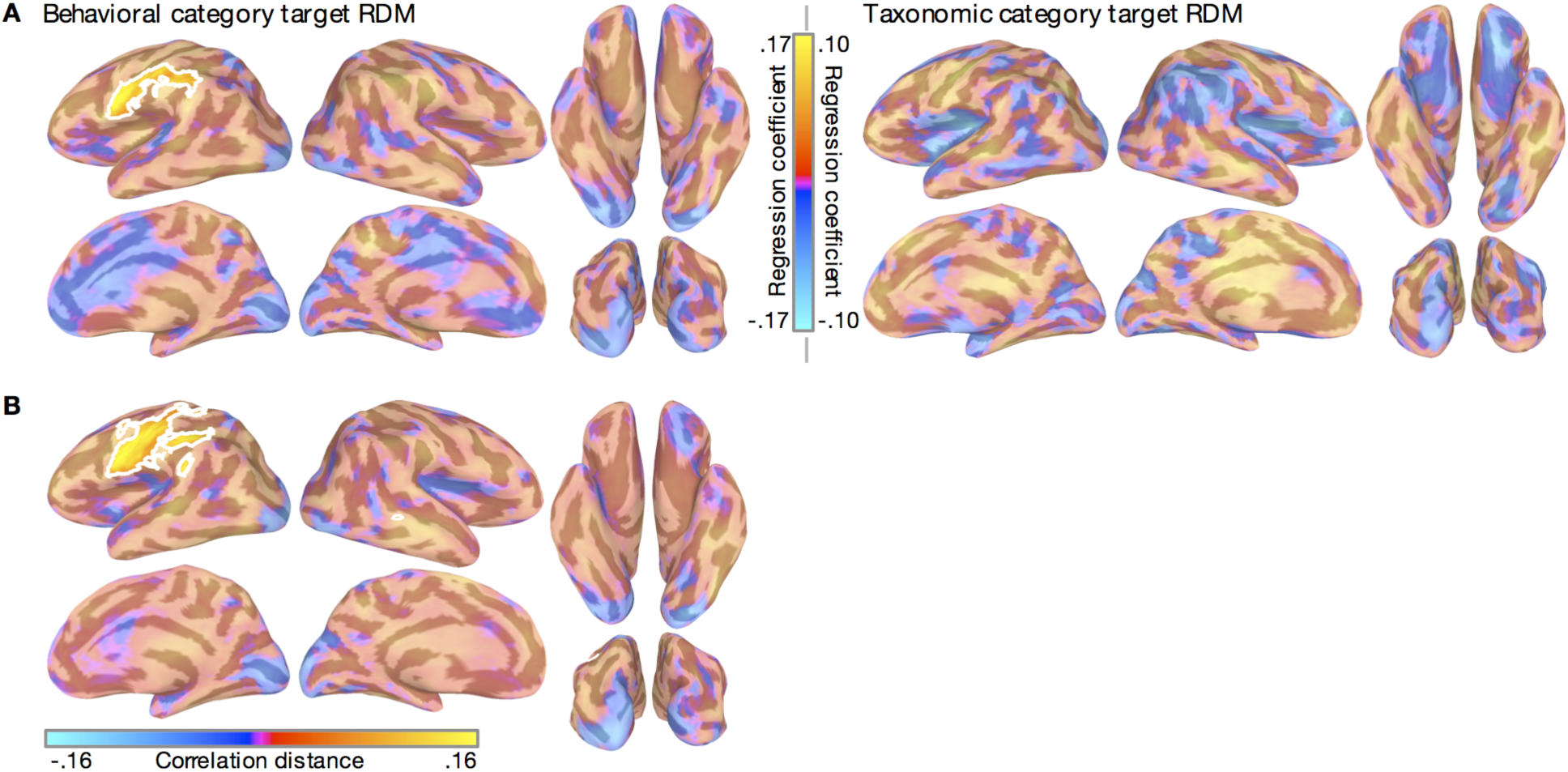
Task differences in searchlight representational geometry. (*A*) Attention-related differences in standardized rank regression coefficients were computed for both the behavioral category and taxonomic category target RDMs. Warm colors represent attentional enhancement for the corresponding semantic information. The range of values on the color bar reflects the mean difference in the regression coefficient. (*B*) Cells of the searchlight RDMs capturing within-category distances for both animal behavior and taxonomy were isolated (see Fig. 4) and tested for attentional enhancement of correlation distance. The absolute values of the within-behavior and within-taxon distances were averaged for each searchlight to compute an index of overall task difference in within-category correlation distances. Clusters surviving TFCE-based correction for multiple comparisons at *p* = .05 (two-tailed test) are displayed at full opacity and outlined with a white contour, while searchlights not surviving TFCE are displayed as partially transparent. TFCE maps were estimated using a Monte Carlo simulation randomly flipping the attention task label. Note that the trend towards an effect of attention to taxonomy in VT cortex on correlation with the taxonomic RDM was not significant in this searchlight analysis but was strongly significant in the ROI analysis that used larger regions. Searchlights in this case included only 100 voxels and cannot capture the more distributed effects observed in the ROI analysis. Furthermore, searchlight analyses are subjected to conservative multiple comparisons correction because of the large number of searchlights.

**Supplementary Figure 5.**
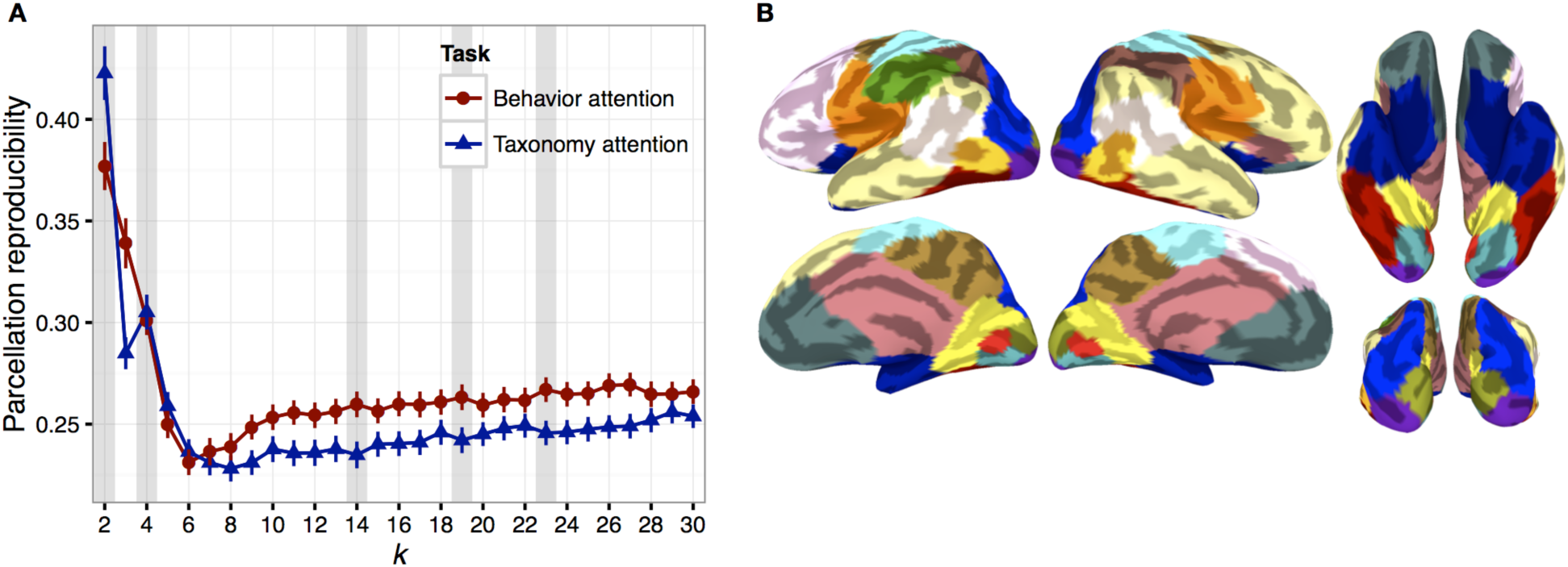
Functional parcellation of the cerebral cortex based on representational geometry. (*A*) Parcellation reproducibility was evaluated using split-half resampling across participants (100 partitions per *k*) separately for each attention task. The mean AMI across the 100 partitions is plotted across the values of *k*, with error bars indicating the standard error of the mean across partitions. Vertical gray bars indicate several local maxima spanning the range of *k* tested. Parcellations at these reproducible values of *k* are visualized on the cortical surface in Supplementary Fig. 5. (*B*) Full parcellation at *k* = 19 for the behavior attention task data. Ten parcels from this solution corresponding to the dorsal and ventral visual pathways were further interrogated in the ROI analysis.

**Supplementary Figure 6.**
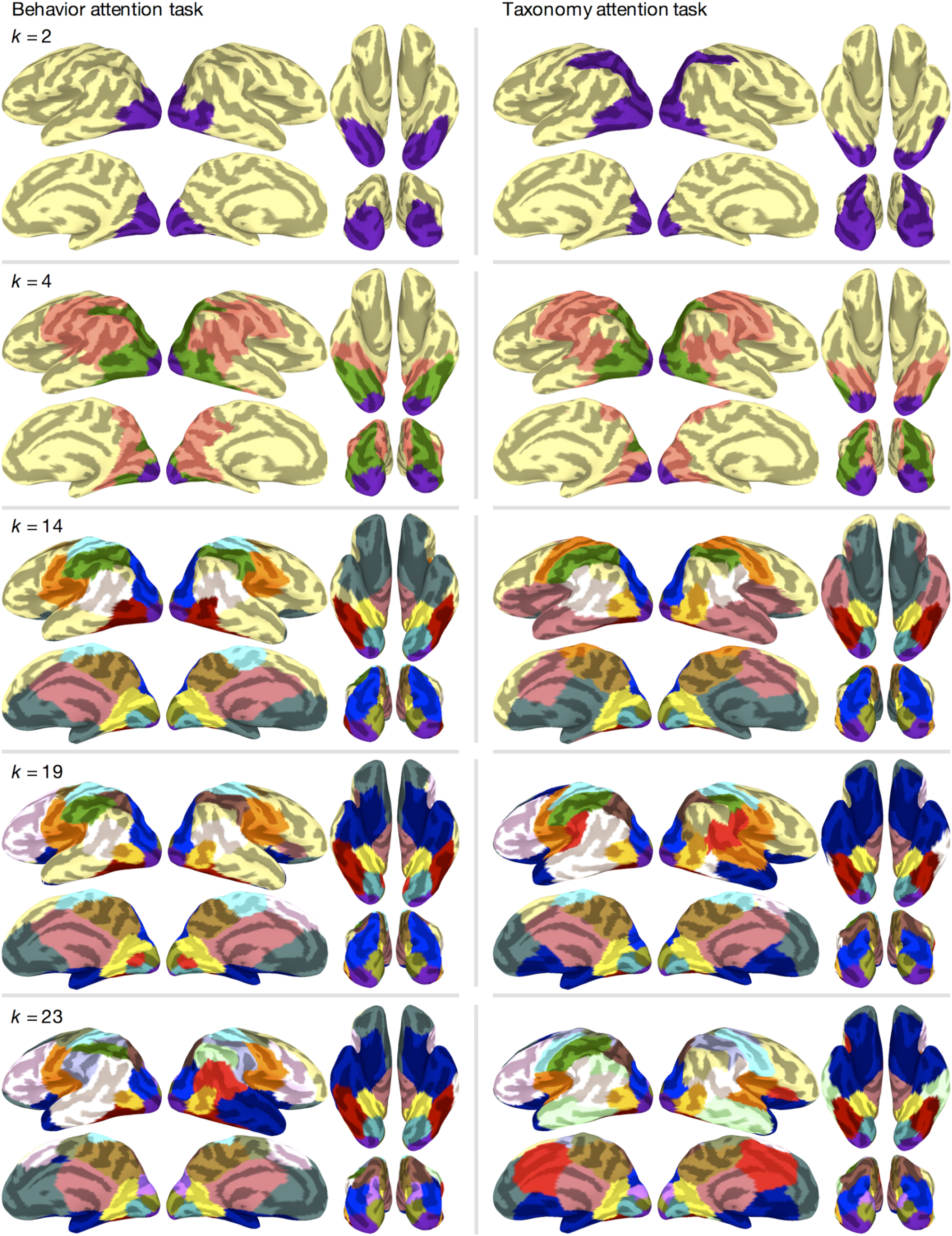
Functional parcellations at reproducible values of *k* for both attention tasks. Parcellation reproducibility was assessed using split-half resampling across participants, and parcellations are depicted for local maxima in parcellation reproducibility (*k* = 2, 4, 14, 19, and 23; corresponding to vertical gray bars in Supplementary Fig. 4*A*). The left column depicts parcellations based on searchlight representational geometries from the behavior attention task and the right column depicts parcellations based on searchlight representational geometries from the taxonomy attention task. The parcellation for the behavior attention task data (left) at *k* = 19 was used for subsequent ROI analysis and is reproduced in Fig. 3 and Supplementary Fig. 4*B*. Colors were assigned manually to avoid similar colors at anatomically adjacent parcels, and to emphasize similar parcels across tasks and values of *k*.

**Supplementary Figure 7.**
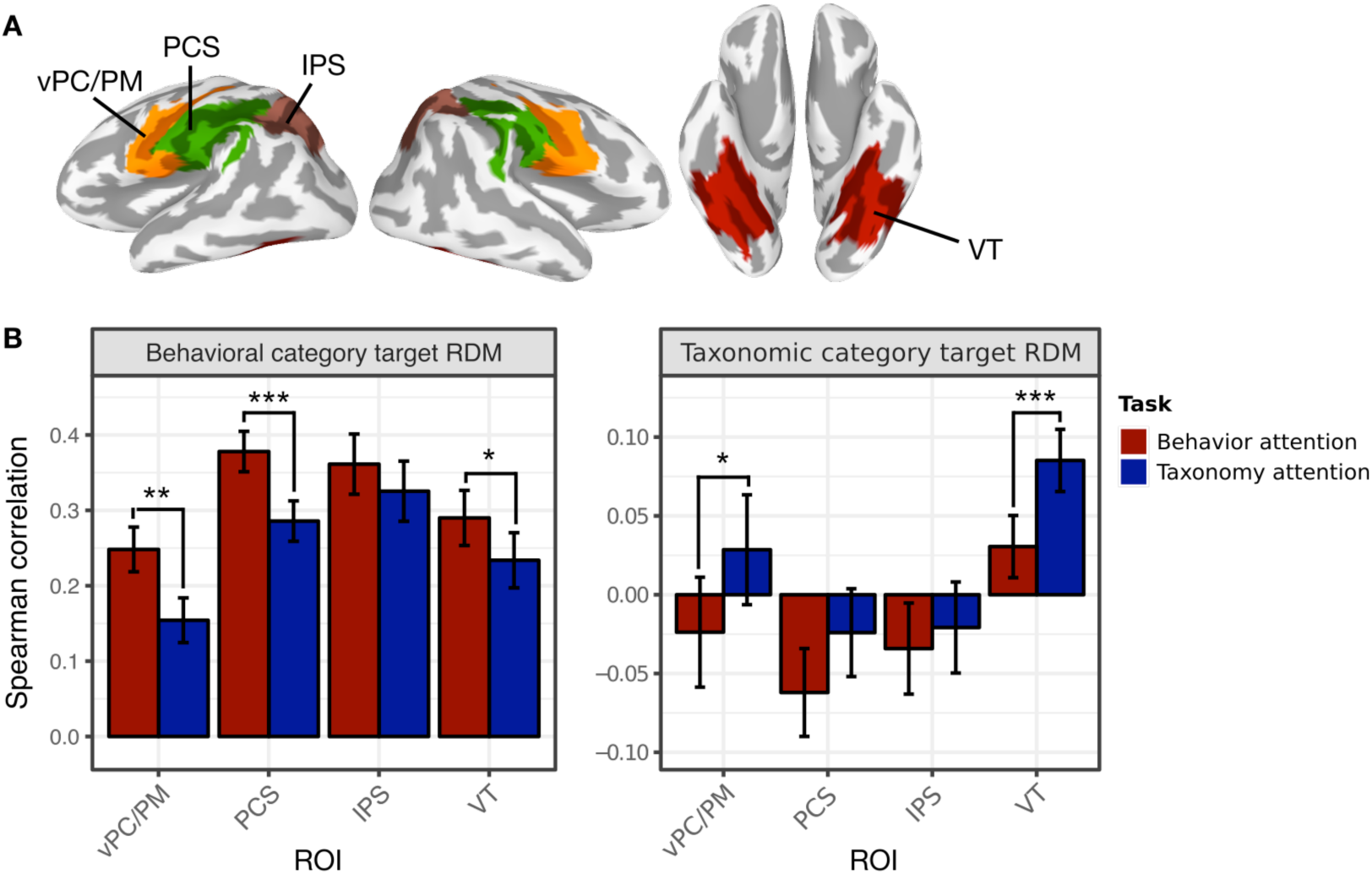
Attention alters representational geometry in anatomically-defined ROIs. (*A*) The following anatomically-defined analogues of four key ROIs were extracted from the FreeSurfer cortical surface parcellation (Destrieux et al. 2010). VT: bilateral fusiform gyri (lateral occipitotemporal gyri), collateral sulci (medial occipitotemporal sulci) and lingual sulci, and lateral occipitotemporal sulci; IPS: bilateral intraparietal sulci, transverse parietal sulci, and superior parietal lobules; PCS: bilateral postcentral gyri, postcentral sulci, and supramarginal gyri extending superiorly to *z* = 50; vPC/PM: bilateral precentral gyri, central sulci, and subcentral gyri (central opercula) extending superiorly to *z* = 50. (*B*) Attending to animal behavior increased Spearman correlations between the observed neural RDM and the behavioral category target RDM in vPC/PM (*p* = .001), PCS (*p* < .001), and VT (*p* = .030). Attending to animal taxonomy increased correlations between the observed neural RDM and the taxonomic category target RDM in vPC/PM (*p* = .043) and VT (*p* < .001). Error bars indicate bootstrapped 95% confidence intervals for within-participants task differences (bootstrapped at the participant level). \**p* < .05, \*\**p* < .01, \*\*\**p* < .001, two-sided nonparametric randomization test.

**Supplementary Figure 8.**
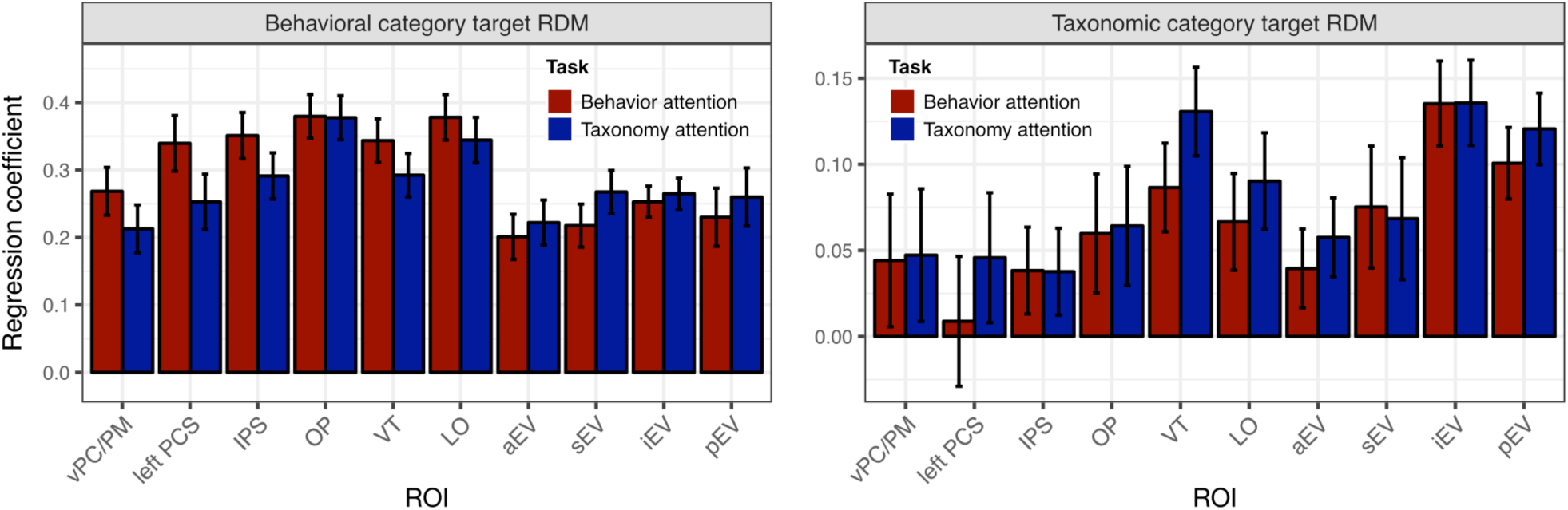
Representational similarity analysis using standardized rank regression. Neural representational geometry in each ROI was modeled as a weighted sum of the behavioral category and taxonomic category target RDMs. Mean regression coefficients for the behavioral category target RDM and taxonomic category target RDM are plotted for both task condition. Error bars indicated bootstrapped 95% confidence intervals for within-participants task differences (bootstrapped at the participant level). This method is identical to the multiple regression used in the searchlight analysis (Figure 2), and reflects qualitatively similar results to the simpler approach using Spearman correlation reported in Figure 3.

**Supplementary Figure 9.**
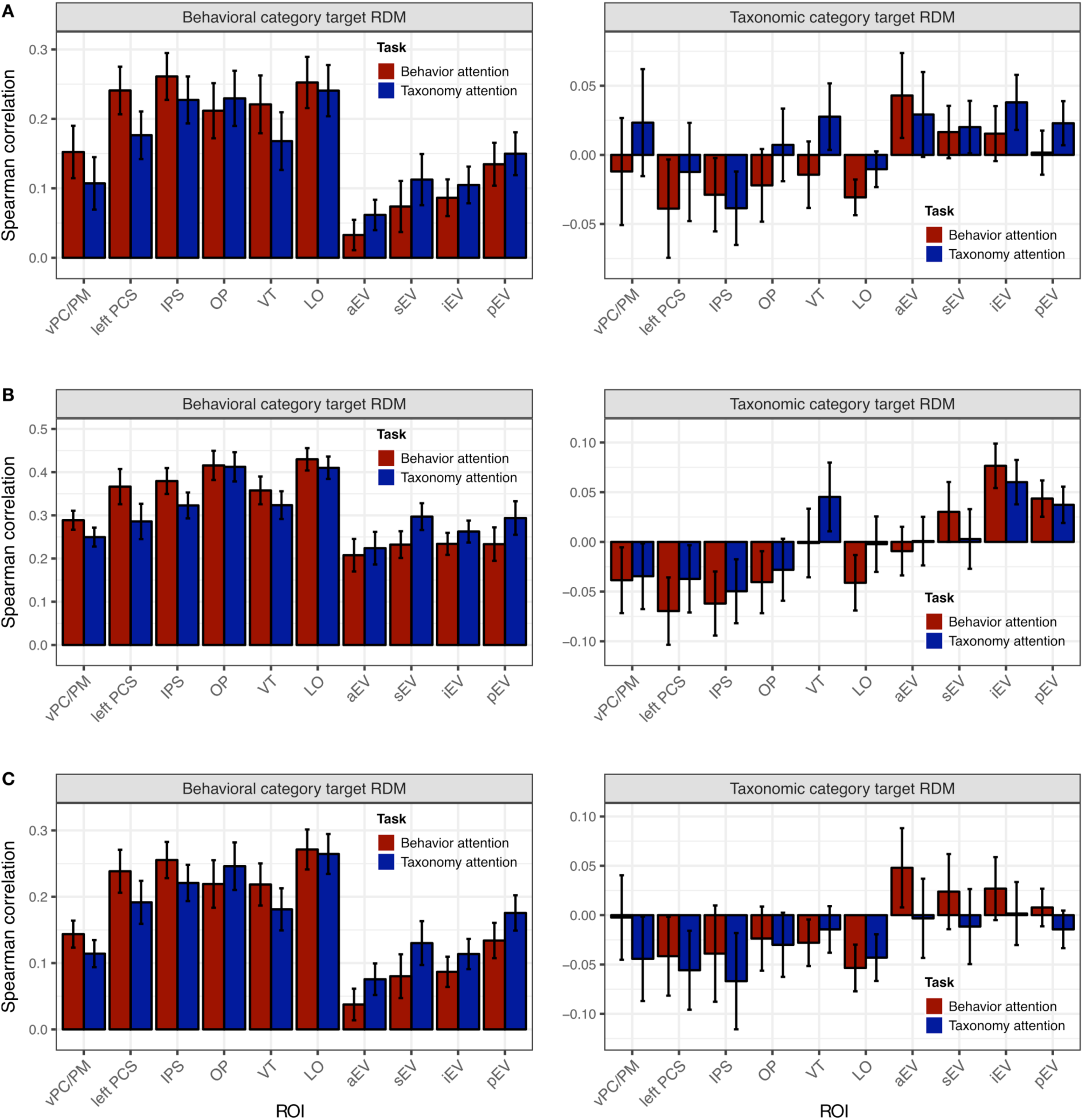
Representational similarity analysis using alternative distance metrics and cross-validation schemes (cf. Figure 3). Neural RDMs for each ROI are compared to the categorical target RDMs using Spearman correlation. All error bars indicate bootstrapped 95% confidence intervals for within-participants task differences (bootstrapped at the participant level). (*A*) Neural RDMs were constructed using Euclidean distance to compute the pairwise dissimilarities between response patterns (rather than correlation distance as in Figure 3). Response patterns were estimated for all five scanning runs for each attention task and neural RDMs were not computed in a cross-validation fashion (as in Figure 3). (*B*) Neural RDMs were constructed using leave-one-run-out cross-validation and correlation distance. Response patterns were estimated separately for each scanning run. For each cross-validation fold, response patterns for four runs were averaged, and pairwise correlation distances were computed between conditions in the averaged runs and the left-out fifth run (for each attention task). This results in a neural RDM with a nonzero diagonal. (*C*) Neural RDMs were constructed using the same leave-one-run-out cross-validation scheme, but using Euclidean distance as the pairwise dissimilarity metric.

**Supplementary Table 1.** Descriptions of video clip stimuli and condition assignments. Each of the 40 video clip exemplars is briefly described. The condition assignments are indicated for each clip. There were two exemplar clips for each condition.

| Description | Behavioral category | Taxonomic category |
| --- | --- | --- |
| Chimpanzee eating a fruit | Eating | Primate |
| Howler monkey eating leaves | Eating | Primate |
| Llama eating cactus fruits | Eating | Ungulate |
| Reindeer grazing on grass | Eating | Ungulate |
| Lammergeier eating carrion | Eating | Bird |
| Hummingbird drinking from flower | Eating | Bird |
| Chameleon eating grasshopper | Eating | Reptile |
| Komodo dragon eating carcass | Eating | Reptile |
| Caterpillar eating its own eggshell | Eating | Insect |
| Ladybug eating mites | Eating | Insect |
| Baboons fighting on rocks | Fighting | Primate |
| Geladas fighting amongst herd | Fighting | Primate |
| Bison butting heads on prairie | Fighting | Ungulate |
| Ibex locking horns on mountainside | Fighting | Ungulate |
| Seabirds fighting on rocks | Fighting | Bird |
| Vultures fighting in the snow | Fighting | Bird |
| Chameleon biting another chameleon | Fighting | Reptile |
| Komodo dragons fighting | Fighting | Reptile |
| Ant and ladybug fighting | Fighting | Insect |
| Stag beetles locking mandibles | Fighting | Insect |
| Baboon running toward water | Running | Primate |
| Monkey running away through tall grass | Running | Primate |
| Juvenile ibex running down mountainside | Running | Ungulate |
| Topi running through herd | Running | Ungulate |
| Penguin running across meadow | Running | Bird |
| Seagull running through cloud of insects | Running | Bird |
| Komodo dragon walking on rocks | Running | Reptile |
| Lizard running across sand | Running | Reptile |
| Ants traveling across sand | Running | Insect |
| Beatle running across dirt | Running | Insect |
| Macaque swimming underwater | Swimming | Primate |
| Snow monkey swimming in hot spring | Swimming | Primate |
| Deer swimming across lake | Swimming | Ungulate |
| Reindeer herd swimming across strait | Swimming | Ungulate |
| Duck swimming across stream | Swimming | Bird |
| Penguin swimming underwater | Swimming | Bird |
| Marine iguana swimming in clear water | Swimming | Reptile |
| Sea turtle swimming near seafloor | Swimming | Reptile |
| Dobsonfly larva swimming toward streambed | Swimming | Insect |
| Water beetle swimming underwater | Swimming | Insect |

**Supplementary Table 2.** Task differences in Spearman correlation for all 19 parcels (Fig. 3). Parcels are listed roughly proceeding from posterior early visual areas anteriorly along the lateral surface, followed by medial structures. Parcel colors reference Supplementary Fig. 4*B*. Extent indicates the number of voxels referenced by all surface-based searchlights in the parcel. The average extent across all 19 parcels was 3,260 voxels (SD = 2,378 voxels). Note that neighboring searchlights overlap spatially and may overlap in the voxels they reference, although these voxels are only counted once for analysis purposes and in each parcel’s extent. Task differences in representational geometry were evaluated by applying (exact) permutation tests to the Fisher transformed Spearman correlations between the observed neural RDM for each parcel and the behavioral category and taxonomic category target RDMs. Reported z-values were derived from the *p*-values returned by the nonparametric randomization test. Negative values indicate decreased Spearman correlation with a target RDM when attending to the corresponding semantic information. Parcel label abbreviations are as follows. pEV: bilateral posterior early visual cortex comprising the occipital pole and posterior lateral occipital sulcus; iEV: bilateral inferior early visual cortex extending from the inferior bank of the posterior calcarine sulcus across the posterior lingual gyrus and posterior transverse collateral sulcus to the inferior occipital gyrus; sEV: bilateral superior early visual cortex encompassing the posterior calcarine sulcus and posterior cuneus; aEV: bilateral anterior early visual cortex including the anterior calcarine sulcus and a portion of the lingual gyrus; LO: bilateral lateral occipitotemporal cortex including the inferior middle occipital gyrus (and human MT+); VT: bilateral ventral temporal cortex including the fusiform gyrus, inferior temporal gyrus, and lateral occipitotemporal sulcus; OP: bilateral occipitoparietal and posterior parietal cortex extending from the lateral occipital sulcus dorsally to the transverse parietal sulcus; IPS: bilateral anterior intraparietal sulcus including the superior parietal lobule; left PCS: left postcentral sulcus, including the postcentral gyrus, inferior parietal lobule (supramarginal gyrus), and anterior intraparietal sulcus; vPC/PM: bilateral ventral pericentral gyri including the ventral central sulcus, premotor cortex, and extending ventrally to include the subcentral gyrus and posterior insula; dPC: bilateral dorsal pericentral gyri and central sulcus extending medially to the paracentral gyrus and posterior medial frontal gyrus; pSTS: bilateral posterior superior temporal sulcus including the posterior middle temporal gyrus and superior temporal gyrus; left dlPFC: left dorsolateral prefrontal cortex extending from the superior frontal gyrus ventrally to the inferior frontal gyrus and extending dorsomedially to the middle anterior medial superior frontal cortex; right dlPFC: right dorsolateral prefrontal cortex extending from the superior frontal gyrus ventrally to inferior frontal gyrus and extending dorsomedially to middle-anterior medial superior frontal cortex, as well as bilateral anterior superior temporal sulcus (aSTS) and middle temporal gyrus, and bilateral temporoparietal junction (TPJ), including the inferior parietal lobule, supramarginal gyrus, and angular gyrus; mV: bilateral medial visual cortex extending from the parietooccipital sulcus across the anterior calcarine sulcus to the parahippocampal gyrus and medial aspect of the fusiform gyrus; Precuneus: bilateral precuneus including subparietal cortex and the marginal ramus of the cingulate sulcus, as well as the bilateral posterior superior frontal sulcus; Cingulate: bilateral middle cingulate cortex, medial subcortical structures, and the right anterior insula; OFC: bilateral orbitofrontal cortex extending posteriorly to include bilateral anterior temporal lobes (ATL; parahippocampal gyrus and temporal pole); mPFC: bilateral medial prefrontal cortex including the anterior cingulate and superior frontal gyrus. See Supplementary Table 3 for tests computed separately for each bilateral homologue and otherwise anatomically discontiguous parcel. \**p* < .05, \*\**p* < .01, two-sided nonparametric randomization test, uncorrected.

| Parcel | Color | Extent | Task differences in Spearman's $\rho$ (z-value) | |
| --- | --- | --- | --- | --- |
|  |  |  | Behavior RDM | Taxonomy RDM |
| pEV | purple | 1,419 | -0.944 | 0.914 |
| iEV | teal | 1,321 | -0.928 | -0.191 |
| sEV | olive | 1,220 | -2.142* | -0.798 |
| aEV | red | 882 | -0.863 | 0.922 |
| LO | gold | 1,333 | 1.652 | 1.707 |
| VT | maroon | 2,063 | 2.326* | 2.567* |
| OP | blue | 3,570 | 0.372 | 0.223 |
| IPS | copper | 2,638 | 2.535* | 0.770 |
| Left PCS | green | 1,356 | 2.784** | 2.095* |
| vPC/PM | orange | 3,995 | 2.228* | 0.649 |
| dPC | cyan | 4,840 | 2.385* | 2.221* |
| pSTS | white | 2,793 | 1.135 | 0.365 |
| Right dIPFC | light yellow | 11,362 | 1.579 | 0.846 |
| Left dIPFC | violet | 5,199 | 1.856 | 0.739 |
| mV | yellow | 2,671 | -0.211 | 0.434 |
| Precuneus | brown | 4,428 | 1.933 | 1.238 |
| Cingulate | dark pink | 3,166 | 1.731 | 0.909 |
| OFC | navy | 5,611 | 1.656 | 0.849 |
| mPFC | dark gray | 3,334 | 0.190 | 0.425 |

**Supplementary Table 3.** Task differences in Spearman correlation computed separately for each anatomically discontiguous parcel. In many cases, the clustering algorithm returned bilateral homologues as one cluster, while in several cases additional spatially discontiguous regions of the cortical surface were included in a single cluster. We split these discontiguous regions into separate parcels based on the neighborhood structure of the cortical surface mesh, then analyzed each parcel separately using nonparametric randomization tests. The average extent across all discontiguous parcels was 1,394 voxels (SD = 1,026 voxels). Z-values were derived from the *p*-values returned by the randomization test, and negative values indicate decreased Spearman correlation with a target RDM when attending to the corresponding semantic categories. In addition to bilateral homologues, the highly diffuse right dlPFC cluster split into bilateral anterior superior temporal sulcus (aSTS) parcels, bilateral temporoparietal junction (TPJ) parcels, and three small parcels in left orbitofrontal cortex (OFC), right anterior insula (aI), and left precentral gyrus (PreC). The Precuneus cluster included bilateral posterior superior frontal sulcus (pSFS) parcels, and the Cingulate cluster included a portion of the left anterior insula (aI). \**p* < .05, \*\**p* < .01, \*\*\**p* < .005, two-sided nonparametric randomization test, uncorrected.

| Parcel | Hemisphere | Color | Extent | Task differences in Spearman's $\rho$ (z-value) | |
| --- | --- | --- | --- | --- | --- |
|  |  |  |  | Behavioral RDM | Taxonomic RDM |
| pEV | L | purple | 712 | -1.309 | -0.098 |
|  | R | purple | 707 | -0.827 | 0.596 |
| iEV | L | teal | 729 | -0.233 | 0.681 |
|  | R | teal | 552 | -1.414 | -0.777 |
| sEV | L | olive | 654 | -2.235* | -1.162 |
|  | R | olive | 566 | -1.725 | -0.991 |
| aEV | L | red | 470 | 0.191 | 1.917 |
|  | R | red | 412 | -1.339 | -1.285 |
| LO | L | gold | 592 | 2.034* | 1.411 |
|  | R | gold | 741 | 0.061 | 0.771 |
| VT | L | maroon | 1,037 | 2.308* | 2.038* |
|  | R | maroon | 1,026 | 1.929 | 2.001* |
| OP | L | blue | 1,878 | 0.474 | 0.406 |
|  | R | blue | 1,692 | 0.328 | 0.408 |
| IPS | L | copper | 980 | 0.378 | 0.130 |
|  | R | copper | 1,658 | 2.602** | 0.692 |
| Left PCS | L | green | 1,356 | 2.784** | 2.095* |
| vPC/PM | L | orange | 1,953 | 2.354* | 1.135 |
|  | R | orange | 2,042 | 1.454 | 0.125 |
| dPC | L | cyan | 2,616 | 2.074* | 2.160* |
|  | R | cyan | 2,224 | 1.704 | 2.095* |
| pSTS | L | white | 1,300 | 1.921 | 0.283 |
|  | R | white | 1,493 | 0.435 | 0.553 |
| Right dIPFC | R | light yellow | 5,004 | 1.532 | 0.562 |
| aSTS | L | light yellow | 1,726 | 1.048 | 0.853 |
|  | R | light yellow | 2,169 | 1.461 | 0.906 |
| TPJ | L | light yellow | 766 | 1.630 | 0.845 |
|  | R | light yellow | 1,117 | 1.488 | -0.113 |
| Left OFC | L | light yellow | 195 | 0.162 | -0.101 |
| Right al | R | light yellow | 138 | 1.691 | -0.070 |
| Left PreC | L | light yellow | 127 | 1.962* | 2.166* |
| Left dIPFC | L | violet | 5,199 | 1.856 | 0.739 |
| mV | L | yellow | 1,374 | -0.261 | 0.511 |
|  | R | yellow | 1,297 | 0.112 | 0.494 |
| Precuneus | L | brown | 1,440 | 0.831 | 0.742 |
|  | R | brown | 1,449 | 1.126 | 0.851 |
| pSFS | L | brown | 803 | 3.487*** | 1.546 |
|  | R | brown | 653 | 0.465 | 0.062 |
| Cingulate | L | pink | 1,380 | 0.677 | 1.358 |
|  | R | pink | 1,234 | 1.725 | 0.841 |
| Left al | L | pink | 552 | 2.079* | -0.569 |
| OFC | L | navy | 2,275 | 1.393 | 0.905 |
|  | R | navy | 2,083 | 1.550 | 0.578 |
| mPFC | L | dark gray | 1,649 | 0.507 | 0.579 |
|  | R | dark gray | 1,685 | 0.077 | 0.287 |

**Supplementary Table 4.** Task enhancement for within-category correlation distances for all 19 parcels (Fig. 4). Reported z-values were derived from the *p*-values returned by the nonparametric randomization test. Positive values in the “within-behavior” column can be interpreted as either decreased within-behavioral category distances when attending to behavior or an increase in between-taxonomic category distances when attending to taxonomy; similarly, positive values in the “within-taxonomy” column can be interpreted as either decreased within-taxonomic category distances when attending to taxonomy or increased between-behavioral category distances when attending to behavior. Negative values indicate the inverse effect. See Supplementary Table 5 for tests computed separately for each bilateral homologue and otherwise anatomically discontiguous parcel. \**p* < .05, \*\**p* < .01, \*\*\**p* < .005, two-sided nonparametric randomization test, uncorrected.

| Parcel | Task enhancement for within-category distances |  |
| --- | --- | --- |
|  | Within-behavior | Within-taxon |
| pEV | -1.287 | 0.606 |
| iEV | -0.664 | 0.145 |
| sEV | -2.200* | -0.659 |
| aEV | -0.696 | 0.534 |
| LO | 0.899 | 1.586 |
| VT | 2.200* | 2.620** |
| OP | 0.563 | 1.301 |
| IPS | 1.917 | 1.390 |
| Left PCS | 3.097*** | 2.584** |
| vPC/PM | 2.705** | 1.134 |
| dPC | 2.090* | 2.620** |
| pSTS | 0.632 | 0.382 |
| Right dIPFC | 1.770 | 1.113 |
| Left dIPFC | 1.617 | 1.023 |
| mV | -0.452 | 0.669 |
| Precuneus | 1.873 | 1.542 |
| Cingulate | 1.501 | 1.278 |
| OFC | 1.669 | 1.207 |
| mPFC | 0.579 | 0.781 |

**Supplementary Table 5.** Task enhancement for within-category distances computed separately for each anatomically discontiguous parcel. In addition to bilateral homologues, the highly diffuse right dlPFC cluster split into bilateral anterior superior temporal sulcus (aSTS) parcels, bilateral temporoparietal junction (TPJ) parcels, and three small parcels in left orbitofrontal cortex (OFC), right anterior insula (aI), and left precentral gyrus (PreC). The Precuneus cluster included bilateral posterior superior frontal sulcus (pSFS) parcels, and the Cingulate cluster included a portion of the left anterior insula (aI). \**p* < .05, \*\**p* < .01, \*\*\**p* < .005, two-sided nonparametric randomization test, uncorrected.

| Parcel | Hemisphere | Task enhancement for within-category distances |  |
| --- | --- | --- | --- |
|  |  | Within-behavior | Within-taxon |
| pEV | L | -1.601 | -0.391 |
|  | R | -1.036 | 0.323 |
| iEV | L | -0.094 | 0.376 |
|  | R | -0.794 | -0.278 |
| sEV | L | -2.221* | -0.985 |
|  | R | -2.019 | -0.669 |
| aEV | L | 0.141 | 1.385 |
|  | R | -1.001 | -0.994 |
| LO | L | 1.918 | 1.461 |
|  | R | -0.582 | 0.866 |
| VT | L | 2.364* | 2.221* |
|  | R | 1.084 | 2.124* |
| OP | L | 1.301 | 1.230 |
|  | R | -0.341 | 0.948 |
| IPS | L | 0.937 | 1.270 |
|  | R | 1.891 | 0.241 |
| Left PCS | L | 3.097*** | 2.584** |
| vPC/PM | L | 2.848** | 1.626 |
|  | R | 2.186* | 0.377 |
| dPC | L | 1.941 | 2.640** |
|  | R | 1.600 | 2.015* |
| pSTS | L | 1.499 | 0.470 |
|  | R | -0.024 | 0.407 |
| Right dIPFC | R | 1.623 | 0.908 |
| aSTS | L | 1.048 | 0.853 |
|  | R | 1.461 | 0.906 |
| TPJ | L | 1.059 | 1.005 |
|  | R | 1.396 | 1.431 |
| Left OFC | L | 0.292 | -0.115 |
| Right al | R | 1.856 | -0.029 |
| Left PreC | L | 2.048* | 2.243* |
| Left dIPFC | L | 1.617 | 1.023 |
| mV | L | -0.050 | 0.674 |
|  | R | 0.254 | 0.314 |
| Precuneus | L | 0.836 | 0.855 |
|  | R | 1.319 | 1.122 |
| pSFS | L | 3.182*** | 1.280 |
|  | R | 0.997 | 0.446 |
| Cingulate | L | 0.602 | 1.430 |
|  | R | 1.538 | 1.144 |
| Left al | L | 1.895 | -0.036 |
| OFC | L | 1.450 | 1.218 |
|  | R | 1.546 | 1.033 |
| mPFC | L | 0.642 | 0.781 |
|  | R | 0.493 | 0.728 |

